# Structural Assessment of the Full-Length Wild-Type Tumor Suppressor Protein p53 by Mass Spectrometry-Guided Computational Modeling

**DOI:** 10.1101/2022.11.11.516092

**Authors:** Alessio Di Ianni, Christian Tüting, Marc Kipping, Christian H. Ihling, Janett Köppen, Claudio Iacobucci, Christian Arlt, Panagiotis L. Kastritis, Andrea Sinz

## Abstract

The tetrameric tumor suppressor p53 represents a great challenge for 3D-structural analysis due to its high degree of intrinsic disorder (ca. 40%). We aim to shed light on the structural and functional roles of p53’s C-terminal region in full-length, wild-type human p53 tetramer and their importance for DNA binding. For this, we employed complementary techniques of structural mass spectrometry (MS) in an integrated approach with AI-based computational modeling. Our results show no major conformational differences in p53 between DNA-bound and DNA-free states, but reveal a substantial compaction of p53’s C-terminal region. This supports the proposed mechanism of unspecific DNA binding to the C-terminal region of p53 prior to transcription initiation by specific DNA binding to the core domain of p53. The synergies between complementary structural MS techniques and computational modeling as pursued in our integrative approach is envisioned to serve as general strategy for studying intrinsically disordered proteins (IDPs) and intrinsically disordered region (IDRs).

## Introduction

The tumor suppressor p53, which is often referred to as the “guardian of the genome”, plays an important role in DNA repair, cell cycle control, and apoptosis in human cells (1-3). Human p53 is biologically active as a homotetramer formed by association of initial dimers (4). It is one of the most studied and prominent representatives of intrinsically disordered proteins (IDPs) (5). The structural and functional characterization of IDPs has been recognized as a crucial task in structural biology, considering the abundance of proteins containing intrinsically disordered regions (IDRs) in all domains of life (archaea, bacteria, eukaryotes) and in viruses (6, 7). More than 40 years after the discovery of the prominent tumor suppressor protein the structure and spatial arrangement of the tetrameric p53 complex, remain elusive. The protein has a modular domain organization (Figure 1) with two ordered domains, the DNA binding domain (DBD) and tetramerization domain (TET), and contains approximately 40% of IDRs comprising the N-terminal transactivation domain (TAD) and proline-rich region (PRR) as well as the C-terminal domain (CTD) (8). The p53 tetramer cooperatively binds to its target duplex DNA in a sequence-specific manner via its DBD (9, 10). The target binding sites consist of two palindromic decameric motifs (half-sites) of the general sequence 5’-RRRCWWGYYY-3’ (R = A, G; W = A, T; Y = C, T) (11). P53 specifically recognizes genes which contain these motifs and regulate their expression levels. Structures of the DBD (12, 13) and the TET (14, 15) are available, as well as combinations of the two (16, 17), but detailed structural information on the N- and C-terminal regions in context of full-length, wild-type human p53 is still missing. Recently, a cryo-electron microscopy (cryo-EM) structure of p53 with the nucleosome, H2A, H2B, H3.1, and H4 plus DNA fragment containing the p53 binding site was proposed, but the low resolution obtained for p53 did not allow resolving its structure in the cryo-EM map (18). Different models have been proposed for the topology and overall conformation of the p53 tetramer in the absence and presence of DNA (19-26). Those models were generated using both experimental as well as computational approaches and differ substantially in their domain organization (Supporting Information, Figure S1).

**Figure 1:**
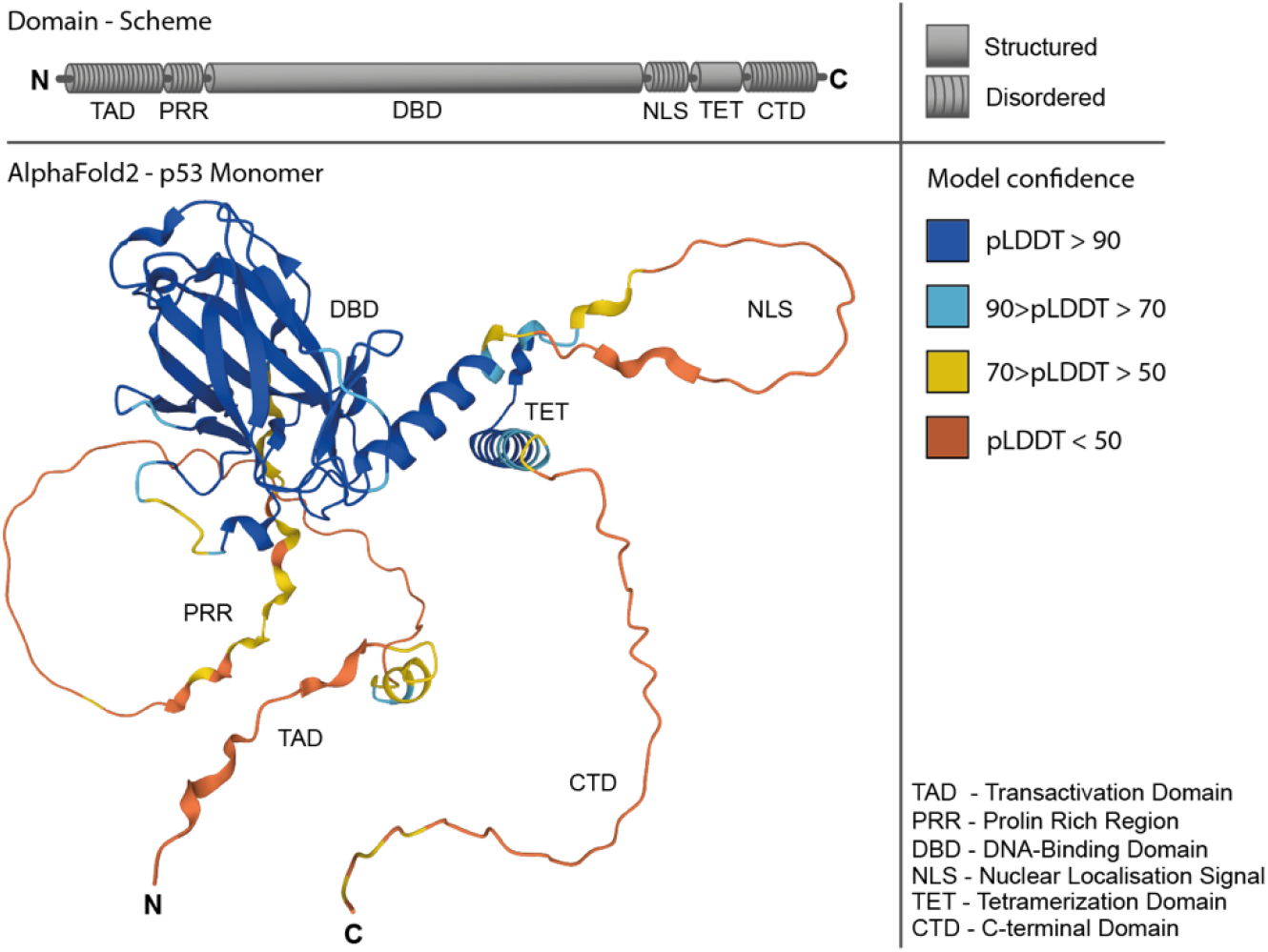
Domain scheme and AlphaFold2 prediction of full-length, wild-type human p53 monomer (cellular tumor antigen p53, https://alphafold.ebi.ac.uk) (32, 67). Upper panel: Structured and disordered regions of p53 are indicated in the domain scheme. Lower panel: AlphaFold2 prediction gives low confidence scores (plDDT) for the IDRs of p53.

Currently, the method of choice for 3D-structure analysis of IDPs is NMR spectroscopy, but complementary structural biology techniques are under continuous improvement (27). These include cryo-EM in combination with structural mass spectrometry (MS) techniques, such as native MS (28), ion-mobility MS (IM-MS) (29), hydrogen/deuterium exchange MS (HDX-MS) (30) and cross-linking MS (XL-MS) (31). A direct computational prediction of IDRs is currently not feasible – if not impossible. With the recent advents of AI-based prediction tools, such as AlphaFold2 (32), it is now possible to generate structures of well-folded protein domains as a starting point, while the prediction of IDRs/IDPs remains a daunting task (Figure 1).

To further investigate the structural properties of p53, with a special focus on the understudied disordered C-terminal region, we adopted an integrative structural biology approach. Specifically, we investigate the topology of p53 in the absence and presence of DNA in solution. In contrast to the majority of structural data published for p53 where IDRs are missing completely or where IDR-derived peptides are used as substitutes (33-35), we employ full-length, wild-type human p53 (36). Our approach combines complementary structural MS methods, AI-based structure prediction with AlphaFold2 (32, 37), data-driven protein modeling, and in-depth scoring and validation of derived models. By this, we gained novel insights into the C-terminal IDR of p53 and evaluate potential structural changes thereof upon DNA binding.

## Experimental

### Chemicals

All chemicals used in this study were of the highest purity available and were obtained from Sigma. Bis(sulfosuccinimidyl)glutarate (BS^2^G) cross-linkers (D_0_ and D_4_) were obtained from Thermo Fisher Scientific; trypsin (porcine, sequencing grade) was obtained from Promega. AspN (purified from *Flavobacterium menigosepticum*, sequencing grade) was obtained from New England Biolabs. DNA response element (top strand: 5’-CGCGGACATGTCCGGACATGTCCCGC-3’ and complementary bottom strand: 5’-GCGGGACATGTCCGGACATGTCCGCG-3’) was purchased from Microsynth. Nano-HPLC solvents were LC/MS grade (HiPerSolv CHROMANORM, VWR).

### Expression and Purification of p53

The HLT-p53 fusion protein was expressed in *Escherichia coli* BL21 (DE3) cells as previously described (36, 38). Cells were harvested after incubation (16 h, 18°C) and lysed by ultrasonication in 50 mM 2-[4-(2-hydroxyethyl)piperazine-1-yl]ethanesulfonic acid (HEPES), 300 mM NaCl, 2.5 mM tris(2-carboxyethyl)phosphine (TCEP), 20 mM imidazole, pH 8.0. The HLT-p53 fusion protein was purified by affinity chromatography on a FPLC system (ÄKTA Pure, GE Healthcare) using a 1-mL HisTrap FF column (GE Healthcare), followed by size-exclusion chromatography (SEC, Superdex 200 column, GE Healthcare) in 50 mM HEPES, 300 mM NaCl, 2.5 mM TCEP, 10% (v/v) glycerol, pH 7.2. After TEV cleavage o/n at 4°C the size exclusion purification step was repeated. The functionality of purified p53 was verified by native MS showing tetramerization of p53 and by a DNA binding assay as shown previously (36, 38).

### Cross-linking

Solutions containing 10 μM wild-type p53 in 50 mM HEPES, 300 mM NaCl, and 2.5 mM TCEP, 10% (v/v) glycerol, pH 7.2, were incubated overnight with or without DNA response element (RE) and cross-linked at 4°C for 1 h with 0.5 mM BS^2^G-D_0_ and BS^2^G-D_4_, respectively. Cross-linkers were dissolved in neat dimethyl sulfoxide (DMSO) immediately before adding it to the protein solutions. The cross-linking reactions were performed in triplicates for each condition and repeated with reversed deuterium labeling. The reactions were quenched by adding ammonium bicarbonate to a final concentration of 20 mM. For XL-MS, samples with and without RE were mixed (1:1) and separated by SDS-PAGE (10% resolving/5% stacking gel). The monomeric p53 band was excised from the gel (Supporting Information, Figure S2) and *in-gel* digested with AspN (37°C, overnight) and trypsin (37°C, 4 h). Peptides were recovered using an extraction buffer (5% (v/v) TFA/acetonitrile; 1:2) and the solution was subjected to LC-MS/MS analysis.

### Mass Spectrometry

Peptide mixtures were analyzed by LC-MS/MS on an UltiMate 3000 RSLC nano-HPLC system (Thermo Fisher Scientifc) coupled to an Orbitrap Fusion Tribrid mass spectrometer equipped with an EASYSpray ion source (Thermo Fisher Scientifc). Peptides were trapped on a C18 precolumn (Acclaim PepMap 100, 300 μm×5 mm, 5 μm, 100 Å, Thermo Fisher Scientific) and separated on a µPAC C18, 50-cm column (PharmaFluidics). After trapping, peptides were eluted by a concave 90-min water-acetonitrile gradient from 3% to 30% B (solvent B: 80% (v/v) acetonitrile, 0.1% (v/v) formic acid) at a flow rate of 300 nl/min. Data were acquired in data-dependent MS/MS mode using stepped higher-energy collision-induced dissociation (HCD) at normalized collision energies of 27%, 30%, and 33%. High-resolution full scans (*m/z* 300 to 1700, R = 120,000 at *m/z* 200) were followed by high-resolution product ion scans (R = 15,000) for 5 s, starting with the most intense signal in the full-scan mass spectrum. Precursor ions with charge states >2+ and <8+ were selected for fragmentation; the isolation window was set to 2 Th. Dynamic exclusion of 60 s (window 2 ppm) was enabled to allow the detection of less abundant ions. Data acquisition was controlled via XCalibur 4.3 (Thermo Fisher Scientific).

### Data Analysis

Cross-links were manually quantified with XCalibur 4.0. Extracted ion chromatograms (XICs) of the precursor masses for BS^2^G-D_0_ and BS^2^G-D_4_ cross-linked peptides were extracted and BS^2^G-D_4_/D_0_ ratios were calculated using the relative intensities of the precursor ions’ isotope distributions. In case of identifying a cross-linked peptide at different charge states or in case it contained an oxidized methionine, ratios were calculated for all possible peptide states. Identification of cross-links (Supporting Information, Figure S3-S16) and “dead-end” cross-links (Supporting Information, Figure S17-S31) was performed with MeroX 2.0.1.7 (39). The following settings were applied: Proteolytic cleavage C-terminally at Lys and Arg (up to 3 missed cleavages were allowed) and N-terminally at Asp and Glu (up to 3 missed cleavages were allowed); peptide length: 5 to 30 amino acids; modifications: alkylation of Cys by iodoacetamide (fixed), oxidation of Met (variable); cross-linker specificity: Lys, Ser, Thr, Tyr, N-terminus; search algorithm: quadratic mode; precursor mass accuracy: 10 ppm; fragment ion mass accuracy: 20 ppm; signal-to-noise ratio > 1.5; precursor mass correction enabled; false discovery rate (FDR) cut-off: 1%, and minimum score cut-off: 20.

“Dead-end” cross-links were analyzed to extract quantitative footprinting data using Mass Spec Studio 2.0 (https://www.msstudio.ca) (40). The following settings were applied: Protein states, plusDNA, minusDNA; reagents, BS^2^G_OH_D_0_ (composition C_5_H_6_O_3_, monoisotopic mass 114.03169), BS^2^G_OH_D_4_ (composition C_5_D_4_H_2_O_3_, monoisotopic mass 118.05680); amino acid modification, methionine oxidation (variable), cysteine carbamidomethylation (fixed); acquisition mode, DDA; fragmentation, HCD; precursor tolerance, 10 ppm; number of tolerable termini, 2; peptide charge, 2-6; peptide length, 5-40; max modifications per peptide, 3; enzyme, trypsin; XIC m/z selector tolerance, 6 ppm; XIC peak to background ratio, 2; fragment mass selector, 20 ppm. Absolute residue labeling yields were normalized to the unbound state. MS data are available via ProteomeXchange with identifier PXD037030.

### HDX-MS

H/D exchange experiments of p53 were performed at room temperature. 1 µl of 10 µM p53 in 50 mM HEPES, 300 mM NaCl, 2.5 mM TCEP, 10% (v/v) glycerol, pH 7.2 was diluted with 9 µl of 10% (v/v) glycerol in 90% (v/v) D_2_O for 30 sec, 1 min or 2 min. Samples were injected to a home-built HDX apparatus, consisting of an HP1200 (Agilent, Santa Clara, USA) HPLC system, cooled to 4°C in a DB951 showcase (Polar Refrigeration, Bristol, UK), and a column setup comprising an Enzymate Protein Pepsin Column, XBridge BEH C18 with VanGuard cartridge 5 mm×2.1 mm, 2.5 µm, 300 Å and an XBridge Peptide BEH C18 column 100 mm×1mm, 3.5 µm, 300 Å (Waters, Milford, USA). Digestion time was 10 min at 25 µl/min 0.1% (v/v) formic acid in water; separation was performed within 6 min at a flow rate of 80 µl/min using a gradient of 3-50% (v/v) acetonitrile in water with 0.1% (v/v) formic acid. Peptic peptides were analyzed with a timsTOF Pro mass spectrometer (Bruker Daltonik, Bremen, Germany) with electrospray ionization source with a source temperature of 50°C, nebulizer gas pressure of 3 bar, dry gas flow of 8 l/min, and capillary voltage of 3500 V.

Mass spectra were recorded in the *m/z* range 100-1700 using the standard proteomics settings of the timsTOF Pro in the COMPASS software. The HDExaminer 3.3.0 software (Sierra Analytics, Modesto, USA) was used to extract the deuterium content of peptic peptides. Data were corrected for back-exchange using data from six independent 10-µl injections of p53 stored in buffer containing 80% (v/v) D_2_O, 10% (v/v) glycerol, and 10% (v/v) H_2_O.

### AlphaFold2 Structure Prediction

A locally installed version of AlphaFold2 (v2.2) was used (32, 37) with the database accession date April 1, 2022. Tetrameric p53 was predicted with the pre-set ‘multimer’ feature (Supporting Information, Figure S32). The amino acid sequence of full-length, wild-type human p53 was used as input.

### Distance Restraints Definition

Experimental distance restraints were defined on the basis of the experimentally obtained BS^2^G cross-linking data. Distance restraints were defined in MODELLER (41) as distance between the Cα atoms of both cross-linked residues within a monomeric p53 chain as *UpperBound* restraint with a mean distance of 15 Å and a standard deviation of 5 Å. For ‘all restraints’, all distance restraints were applied to all four p53 chains symmetrically. For ‘reduced restraints’, 6 out of 12 experimentally determined cross-links were randomly selected for each p53 monomer, resulting in an asymmetric selection of tetrameric p53 structures. For ‘random restraints’, two residues within the amino acid sequence of p53 (residues 301-393) were randomly selected, 12 random restraints were defined and applied to all monomeric p53 chains symmetrically (Supporting Information, Figure S33).

### Model Refinement

The initial AlphaFold2 model of tetrameric full-length p53 was truncated to the region comprising residues 301-393 and used as initial template for the MODELLER refinement. MODELLER was run by default with activated variable target function method (VTFM) and molecular dynamics (MD) optimization. Distance restraints were redefined for each unique refinement as described above. For the ‘all restraints’ condition, 12,000 unique models were predicted. For ‘reduced restraints’ and ‘random restraints’ conditions, 100 unique restraints were defined and 20 unique models were predicted for each condition. In total, 16,000 refined models were predicted (Supporting Information, Figure S33).

### Model Validation and Scoring

All predicted models were analyzed regarding cross-link distance as well as labeling satisfaction. BS^2^G cross-linking data generated in this study as well as published disuccinimidyldibutyric urea (DSBU) cross-linking data (38) were used, Cα-Cα distances were determined, and a scoring matrix was applied (Supporting Information, Figure S34; Supporting Information, Table S1). For satisfaction of the footprinting data, the solvent-accessible surface area (SASA) was calculated with the MSMS-toolkit (42), the pK_a_ values were calculated with PropKa (43). For SASA scoring, a scoring matrix was applied (Supporting Information, Figure S35; Supporting Information, Table S2). The radius of gyration was calculated with Python code, adapted from https://github.com/sarisabban/Rg.

## Results

The tumor suppressor p53 poses immense challenges for deriving structure-function relationships due to its high content of intrinsic disorder (44). The majority of p53 structural studies has been carried out by using truncated p53 variants where the IDRs have been partially or completely removed (33) or by using peptide representatives derived from p53’s IDRs (34, 35). In this work, we use full-length, wild-type human p53 without any truncations or modifications to gain more detailed mechanistic insights into the structure-function relationships of p53, with a special focus on the C-terminal region. The C-terminal region of p53 plays a crucial role in target recognition and is highly modified by acetylation, ubiquitination, phosphorylation, etc. (45), but detailed molecular information on this region is still limited. It has been hypothesized that the lysine-rich C-terminal region of p53 assists in DNA-binding, mainly mediated via electrostatic interactions. Then, p53 slides along the DNA until the response element is recognized by the DBD of p53 for high-affinity binding (46).

### Structural Investigation of Full-Length p53

One cornerstone of our structural investigation is XL-MS as it is one of the few techniques in structure biology that can handle IDPs/IDRs. In this study, we employ the homobifunctional cross-linker BS^2^G that targets primary amine groups in proteins, i.e., lysines and N-termini (47). Its reactivity makes BS^2^G an ideal reagent to specifically target the CTD of p53 with its high lysine content. Additionally, two BS^2^G isotopologs, BS^2^G-D_0_ and BS^2^G-D_4_, are commercially available (Figure 2A). These reagents generate distinct MS patterns, simplifying cross-link identification and quantification (Figure 2A). We identified several unique intramolecular cross-links in p53, mainly covering the disordered NLS and CTD (Figure 2B; Supporting Information, Table S3). BS^2^G cross-links were quantified in the mass spectra to deduce eventual structural changes in the DNA-free versus DNA-bound states of p53 (Figure 2A). Cross-links identified in at least two out of six replicates were used for quantification. Strikingly, the cross-link patterns did not show major differences between DNA-bound and DNA-free states, indicating similar conformational ensembles of p53 (Figure 2C).

**Figure 2:**
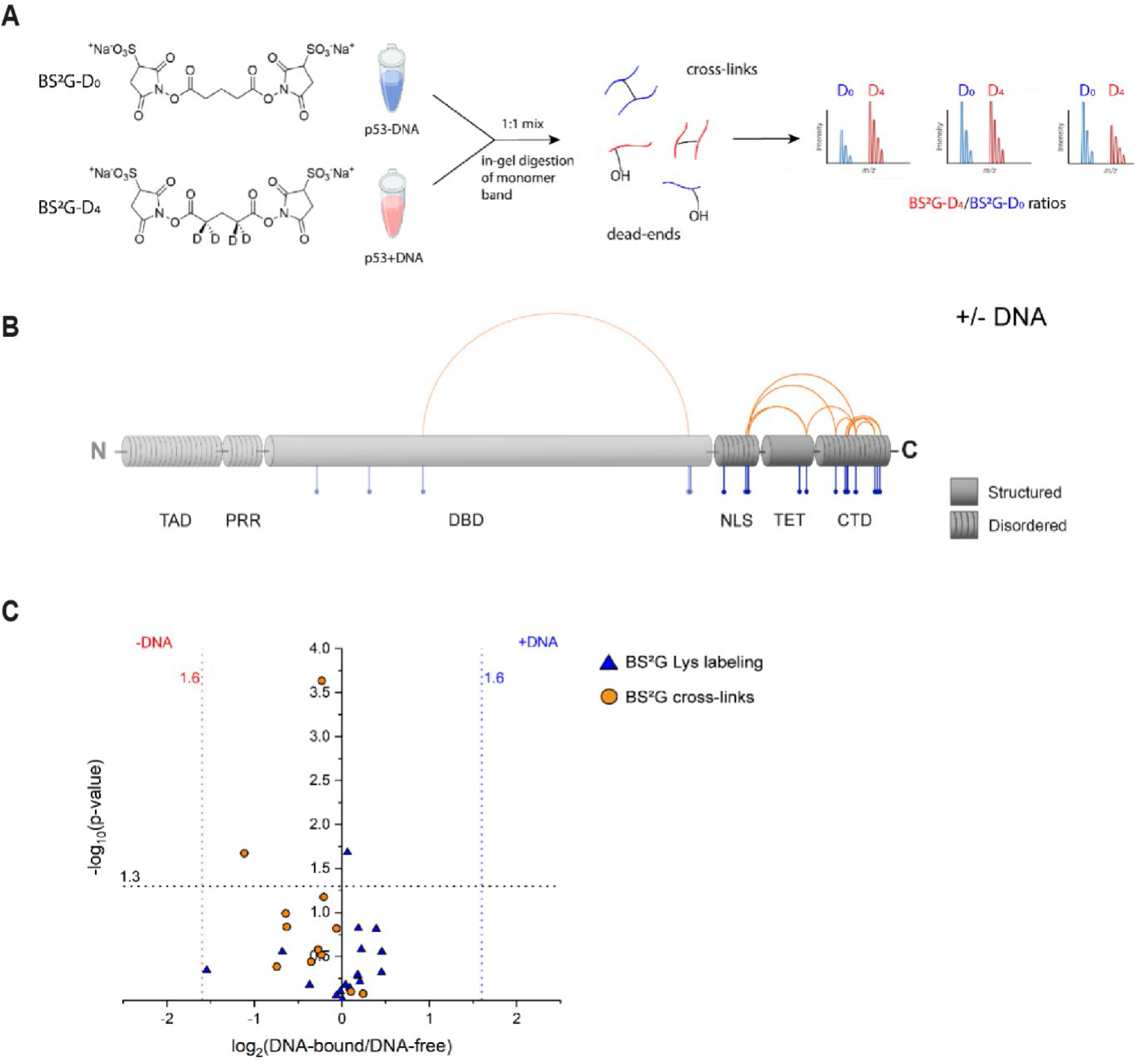
(A) Quantitative cross-linking of full-length p53 was performed with the lysine-reactive, isotope-labeled cross-linker pair BS^2^G-D_0_/D_4_ to reveal differences between DNA-free and DNA-bound p53. (B) Schematic overview of BS^2^G cross-linking (orange) and footprinting data (blue). Structured and disordered regions of p53 are indicated. (C) Comparison of cross-linking and footprinting data for DNA-free (left hand site) and DNA-bound (right hand site) p53. Vertical asymptotes represent the fold change thresholds for determining if a cross-linked or labeled peptide is enriched in the DNA-free or DNA-bound state of p53. The horizontal asymptote defines the statistical significance limit of the data points (p-value < 0.05).

The same p53 samples were used to extract protein footprinting MS data by quantifying “dead-end” cross-links (Figure 2B)(48). “Dead-end” cross-links are peptides that are modified by a partially hydrolyzed cross-linker. These species do not provide direct distance information, but yield important insights into the overall topology of proteins based on solvent accessibilities and local pK_a_ values of specific amino acid side chains. The combined information from interpeptide and “dead-end” cross-links is a highly valuable input for 3D-protein modeling. The mean labeling rates range between ∼2 to 30 %, and residues were classified as buried (< 5% labeling rate), partially accessible (< 10% labeling rate), and accessible (> 10% labeling rate) (Supporting Information, Table S4). Conclusively, the footprinting results indicate that the accessibility of p53 CTD remains unaltered upon DNA binding (Figures 2B and 2C). This finding is in agreement with XL-MS data and suggests that the C-terminus of p53 retains its overall topology upon DNA binding.

### Structural Modeling of p53

The cross-links identified in this study as well as in previous work (38) are predominately located in the C-terminal region of p53, specifically in the TET with flanking IDRs. The data-driven modeling of p53 is therefore focused on the C-terminal region (residues 301 to 393; comprising NLS, TET and CTD). The initial tetrameric full-length p53 model was generated with AlphaFold2 (32, 37) and the C-terminal regions were isolated. The ordered TET shows high confidence and is in agreement with previously solved X-ray and NMR structures (14, 15). However, AlphaFold2 gives only low-confidence models for the disordered regions of p53 (NLS, CTD; Figure 1; Supporting Information, Figure S32). To optimize the structures by satisfying the distance restraints, the initial AlphaFold2 models were subjected to MODELLER. The experimentally derived distance information as well as a reduced and a randomized data set (see Materials and Methods for details) were used to define distance restraints and 16,000 p53 tetrameric models were generated. The refinement strategy is shown in Figure 3A.

**Figure 3:**
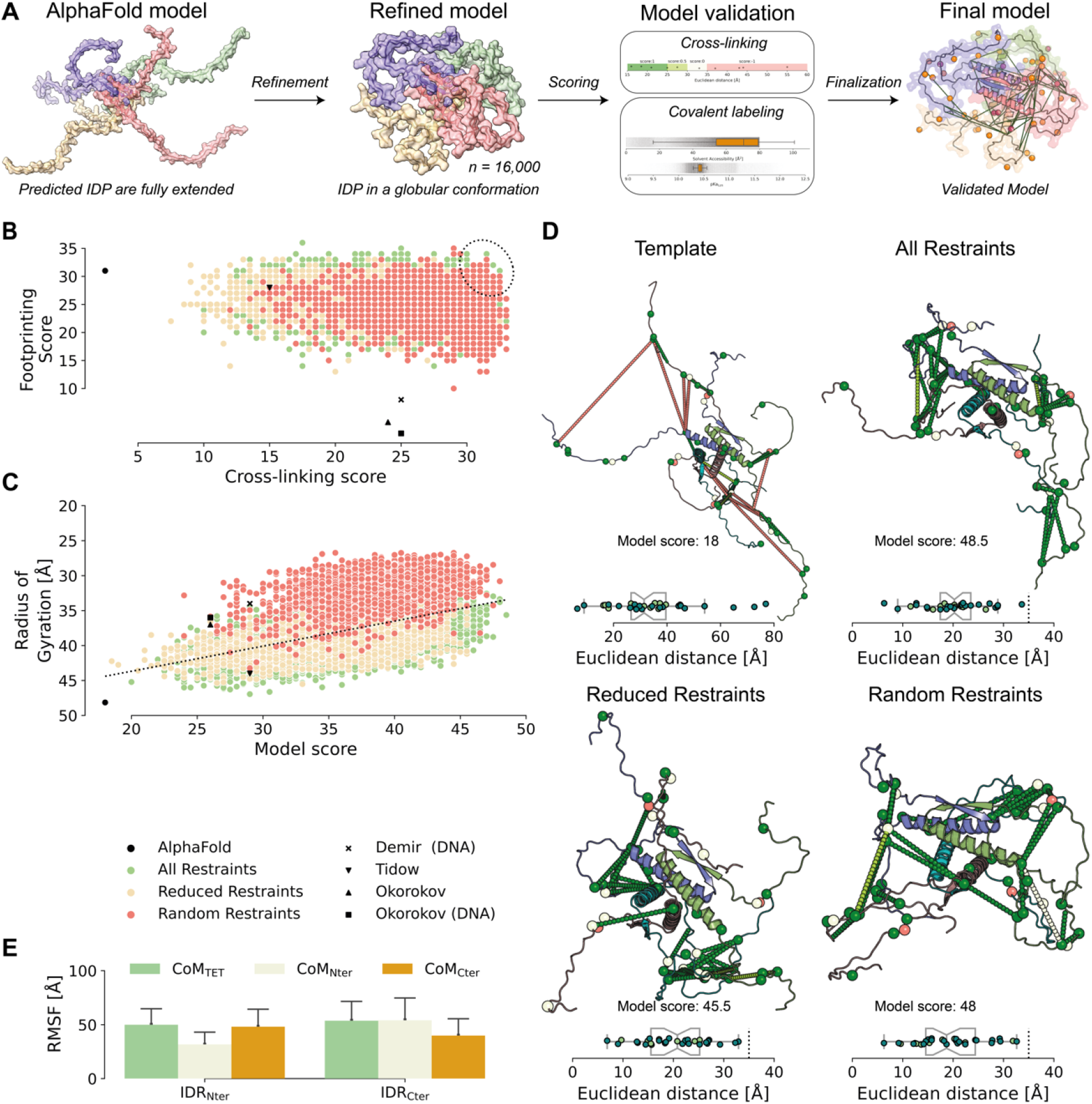
AI-based and knowledge-based remodeling of the disordered C-terminal region of p53. (A) Refinement workflow: The initial model was generated using AlphaFold2. The initial model was refined using experimentally and randomly generated restraints, and the 16,000 models generated were classified and validated using cross-linking and footprinting data. (B) Distribution of all models generated. The dotted ellipsoid marks the highest ranked models. (C) Variation of the radius of gyration (Rg) as a function of the model score. High compaction (low Rg) correlates with high scores of the models. (D) Presentation of input model (Template) and top-ranked refined models (All Restraints, Reduced Restraints, and Random Restraints). For details see Materials and Methods part. The dotted lines indicate cross-links, the spheres present labeled residues. The box-plots indicate Euclidian distances between the Cα-atoms of cross-linked residues; BS^2^G cross-links (light green) and DSBU cross-links (teal). (E) Structural variation of disordered p53 termini. The center of mass (CoM) of the ordered tetramerization domain (TET), as well as the disordered NLS (Nter)- and CTD (Cter) in each refined model were used to calculate the variations between the refined structures.

### Validation of the Refined p53 Structure

The generated models were validated with the experimentally determined cross-links (BS^2^G, 12 cross-links) in addition to previously published cross-links (DSBU; 21 cross-links) (38), and BS^2^G footprinting data. A stepwise scoring matrix was applied with a Cα-Cα distance threshold of 25 Å for BS^2^G and 30 Å for DSBU. Distances of 10 Å above these values were included to account for the high flexibility in the IDRs, based on the overall distance distribution (Supporting Information, Figure S34). In addition, footprinting data yield information on the labeling efficiency of specific lysine residues, depending on their reactivity and accessibility. The reactivity is encoded by the pK_a_ values of lysines and was calculated for each model with PropKa (43), while the accessibility can be described by the SASA and was calculated with the MSMS-tool (42). To access the distribution of pK_a_ and SASA values of each lysine residue, all models were plotted against the experimentally determined accessibilities of lysines (Supporting Information, Figure S35). Notably, the pK_a_ value does not allow differentiating the results, while the SASA shows a population of lysine residues with reduced accessibility, correlating with footprinting data (Supporting Information, Table S4). For scoring, a stepwise rating matrix was applied that penalizes clear outliers (Supporting Information, Figure S35).

For model selection, XL-MS and footprinting data were plotted against each other, and the highest ranked solution of each refinement strategy was picked (Figure 3B). A clear improvement of scoring, both in cross-link distance and footprinting data satisfaction, is visible when compared to the AlphaFold-generated input model and already published tetrameric p53 models (Figure 3B), which clearly indicates an experimentally-driven optimization of computationally predicted AlphaFold models.

One striking observation is that the refined p53 models appear to be more compact than already existing models. To underpin this, we calculated the radius of gyration (Rg) for each model. There is a significant correlation between the scoring and the Rg of p53 (Figure 3C; Supporting Information, Figure S36). Selecting and aligning the top-ranked models reveal an ordered and structurally preserved TET, but the neighboring IDRs exhibit various conformational states (Figures 3D and 3E).

An identical or similar kind of compaction of the p53 tetramer has already been observed previously (25, 38, 49), but is not reflected in alternative models (Supporting Information, Figure S37) (20). Using our experimental data and our scoring, all previously proposed models of the C-terminal region perform worse in the overall scoring than our compact models (Figure 3C).

We additionally validate our p53 tetrameric models with HDX-MS data. HDX-MS displays protein dynamics and changes in a time-dependent manner and complements XL-MS and footprinting data (30, 50, 51). In HDX, backbone amide protons of a polypeptide chain are limited in their H/D exchange if they are involved in hydrogen bonding networks. We mapped the deuterium incorporation kinetics of full-length p53 tetramer for the first time, yielding fast exchange rates for the IDRs, TAD, NLS, and CTD, and slow exchange rates for regions with known secondary structures, DBD and TET (Figure 4A). Plotting the accessibilities of the backbone amide protons in the AlphaFold2 template as well as in the previously proposed models (Supporting Information, Figure S1, Demir and Tidow models) and top-ranked refined models (Figure 3D), the expected correlation with deuterium incorporation in HDX was confirmed (Figure 4B; Supporting Information, Figure S37).

**Figure 4:**
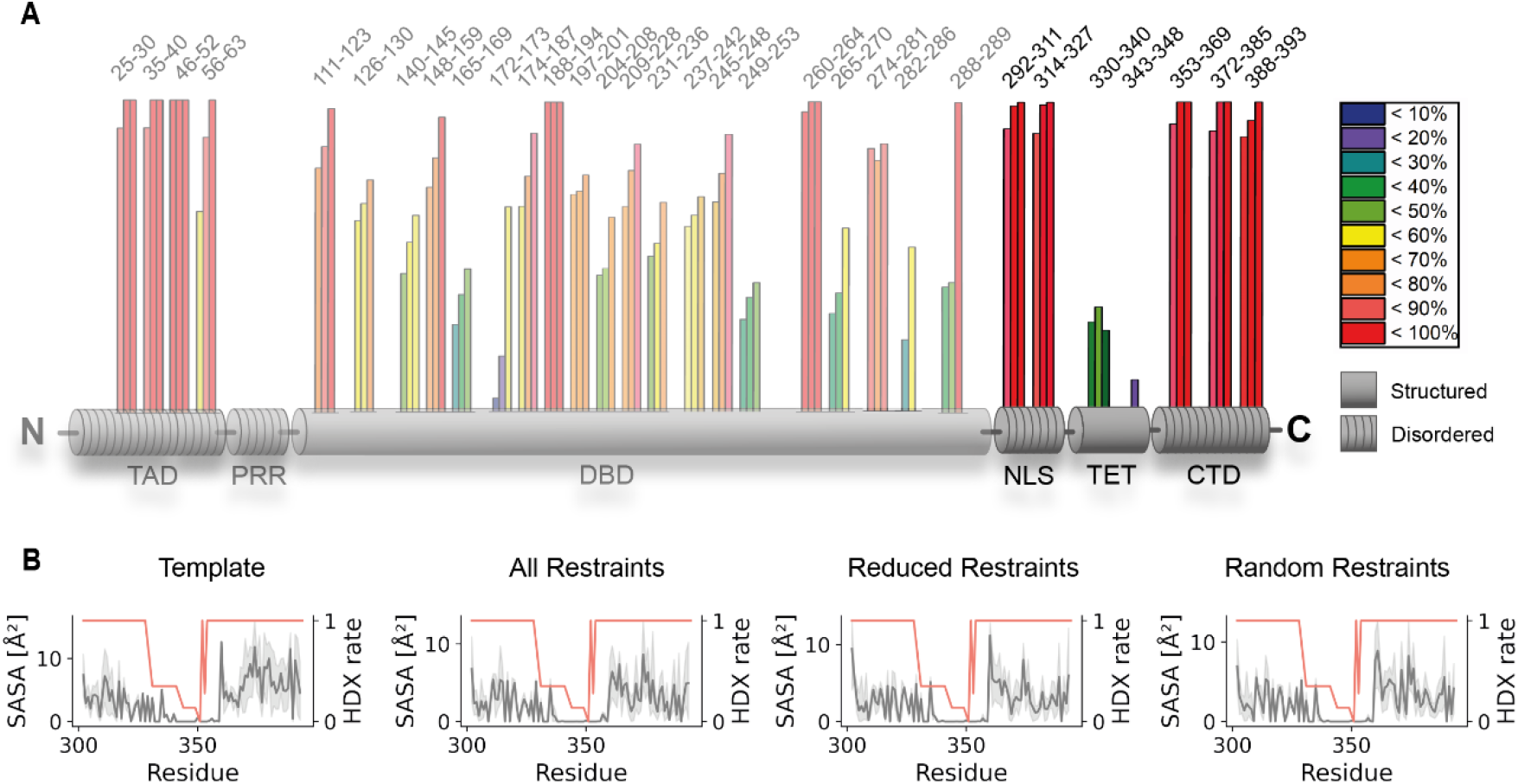
(A) HDX-MS of backbone amide protons in p53. Three bars (HDX after 30 s, 1 min, and 2 min) are presented for each peptide analyzed; amino acid numbers are shown above. Data from NLS, TET, and CTD confirm the top-ranked refined models. (B) Models (Figure 3D) show a high correlation between SASA (grey) and HDX (orange). The disordered and accessible nature of p53’s IDRs is reflected by high SASA and HDX rates, while the structured TET (residues 326-356) exhibits low SASA and HDX rates.

## Discussion

The majority of published p53 structures (12, 15, 16) as well as proposed models (19–21, 25, 26) either lack information about the CTD (residues 363-393) or are inconsistent among each other. This leads to inherent difficulties in interpreting experimental data regarding the CTD, especially in the functional and cellular context of tetrameric p53. By combining complementary techniques of structural MS with AI-based computational modeling we were able to create a conformational ensemble for C-terminal region of tetrameric human wild-type p53(TET, NLS and CTD), satisfying the experimental data.

### Computational Modeling

The default AlphaFold2 model generation includes template identification, structure prediction, and AMBER relaxation (52). The AlphaFold2 relaxation does not have a fixed time scale, but a scoring scale. If the model fulfils certain criteria, MD is stopped. Although this approach works very well in the context of well-structured, globular proteins we encountered challenges by generating models for IDPs with three major key factors. These include (a) lack of template structures due to their disordered nature, (b) MD simulations are performed in explicit water although the choice of force field and the type of water solvation have an impact, and (c) AlphaFold2 is designed for ordered proteins by default. In addition, another important aspect is the hypervariability of IDRs that lead to low-confidence AlphaFold2 models (53). Therefore, a refinement of IDP/IDR models generated with AlphaFold2 that are based on experimental distance restraints is inevitable. In addition to generating distance restraints for computational modeling, protein footprinting adds another layer of validation as only solvent-exposed residues will be labeled efficiently. Notably, local pK_a_ values alone are not suitable for explaining different labeling efficiencies of lysine residues in p53, which is most likely due to the solvent-exposed nature of lysines in the disordered regions. Although these lysines are buried in a subset of derived models they are apparently not involved in ionic interactions or extended hydrogen bonding networks and have therefore similar pK_a_ values. This is reasonable for IDRs as lysine residues that form beforementioned interactions will likely be part of an ordered structure, while buried lysines with unchanged pK_a_ values will most likely be located in an IDR. The final models show a compact shape of p53, which is in agreement with all input data. The disordered character of p53 is recapitulated in the assembly of selected models as no clear conformation is visible and each model has a unique localization of the IDRs (Figures 3D and E). Interestingly, even without similar conformations, the radii of gyration are similar among the highest-ranked models. This could have an implication on how IDRs behave in solution, at least those of p53. As the IDRs are apparently part of the same conformational ensemble, independently of RE-DNA binding, no secondary structure or domain folding was assumed. This scenario might be completely different for IDRs that undergo conformational changes upon binding or during substrate channeling, e.g. in pyruvate dehydrogenase complex (54).

### Compaction of p53 Tetramer

A major finding of our study is the compaction of the whole NLS-TET-CTD arrangement in tetrameric full-length, wild-type human p53, which was initially indicated by our experimental data and is also mirrored in our generated models. The first published structural model by Okorokov *et al* in 2006 (19) was generated by cryo-EM utilizing murine p53, while all other studies were performed with human p53 (55, 56) (Supporting Information, Figures S1B and D). The first human-based, tetrameric p53 structural model derived by the Fersht group (20) differed dramatically in the DNA-free (extended cross-shape, Supporting Information, Figure S1A) and DNA-bound form (compact basket shape, Supporting Information Figure S1C). The DNA-bound tetrameric p53 model was further refined by a follow-up study utilizing negative stain EM (21). Here, different conformational ensembles (classes) were observed, which already gave hints of inter-domain proximities of CTD and NLS, as well as DBD and the DNA itself. Single-molecule FRET (smFRET) experiments from the same study also showed a spatial proximity between the very C-terminus of p53 and the L1-loop (residues 112-124) within the DBD in presence of RE-DNA. However, in the absence of DNA, an equilibrium might exist between the formerly published, extended cross-shaped p53 tetramer (20) and a more compact arrangement as observed for the DNA-bound p53 tetramer (21). The existence of an equilibrium state is controversially discussed due to the limited distance information of smFRET data as only one single labeled construct was used. Also, a spatial rearrangement of the flexible CTD was not discussed, but only a complete rearrangement of the whole p53 tetramer (21). In contrast, our data show that in both cases, DNA-free and DNA-bound states, a more compact p53 structure is present. Another publication agreeing with our findings of a more compact p53 tetramer, both in the DNA-bound and DNA-free states, is based on ion mobility coupled to MS (49). Here, experimentally derived collisional cross sections for DNA-free, full-length p53 and DNA-bound p53 are smaller than the values calculated for the extended model structures (20, 49). Also, collisional cross sections were found to be more similar between both states (+/-DNA) than expected from the models. At that time, the observed compaction was described as a gas phase collapse, which can be excluded in our findings as XL-MS, footprinting, and HDX-MS data were exclusively collected in solution at near physiological buffer conditions. In the most recent MD study of a reconstructed p53 tetramer in complex with RE-DNA a compaction and even a contact of the CTD to the bound DNA molecule is described, supporting our findings (Supporting Information, Figure S1E)(25). Together, all abovementioned p53 structural studies show similar results to our MS-based data and agree with our p53 models.

### Modes of DNA Binding

Initially the CTD was characterized as negative regulator impairing specific DNA binding (8, 57, 58). Later, it was recognized to be a positive regulator being responsible for various functions of the p53 tetramer related to DNA-binding (59, 60). The CTD has been decribed to facilitate p53 binding to long non-specific DNA molecules, as well as to chromatin (61), enabling lateral movement along the DNA via electrostatic interactions. This however does not seem to be entirely true as the CTD might be involved in specific DNA binding as well (62). A two-mode binding model switching between a sliding- and recognition-mode was introduced, termed S- and R-modes (46, 63). This two-mode DNA-binding model suggests a hemi-specific binding, with one dimer of tetrameric p53 binding to DNA in a site-specific manner, while the other p53 dimer binds non-specifically to noncognate DNA transitioning into a fully specific, DNA-bound state upon response element recognition (46). If we reflect our data in the light of that model (S- and R-modes), we obtain a snapshot even for specific DNA binding. We captured a conformational ensemble for one specific 26-bp RE-DNA (one sequence, one length, see also Materials and Methods section) bound to p53, which might reflect the recognition mode (R-mode). A comparison of that two-mode DNA-binding model proposed by Tafvizi *et al* (46) with the EM-based model of Melero *et al* (21) revealed the following: In the latter model, EM maps were grouped into classes that depend not only on the sequence of the RE-DNA, but also on the length of the individual DNA fragment (18-mer *vs* 44-mer *vs* 60-mer). Also, the number of p53 conformational states observed differed depending on the nature of the bound DNA-fragment. In our study, we used an intermediate length (26-mer) RE-DNA with high specificity and affinity towards the DBD (49, 64). This is a strong hint that we indeed captured one of the specific DNA-binding modes, in which the CTD of p53 might be involved to stabilize the R-mode. Our findings are also in line with the reaction kinetics of the cross-linking and labeling reagents used herein thatcapture dynamics on the minutes to seconds time scale. We are therefore confident that our XL-MS setup probably captures a conformational ensemble of full-length p53 bound to a specific RE-DNA in the R-mode.

There are however still discrepancies whether CTD and DBD are both binding to DNA, interdependently of each other, or if the CTD controls site-specific binding of the DBD to the DNA by inducing conformational changes within the DBD itself (26, 62). If there were conformational changes upon DNA-binding on the sub-second time scale, like a channeling of the DNA from the CTD to the DBD, faster reaction kinetics available with photo cross-linkers (diazirines, incorporated photo-amino acids) would be needed (65, 66). To capture the conformational dynamics of the highly complex binding events between full-length p53 and DNA, integrative approaches are inevitable. One technique alone will not be able to grasp the full picture, but structural MS combined with AI-based computational modeling allows getting insights into the high complexity of p53’s conformational ensembles.

## Conclusion and Outlook

As it becomes increasingly obvious that a single study will never be powerful enough to completely unravel the whole mechanistic machinery of p53 tetramerization, DNA-binding, and movement along the bound DNA, integrative approaches are required for a better understanding. We contribute to this tremendous task by delivering a workflow for the in-solution investigation of full-length, wild-type human p53 with a consecutive structural modeling pipeline. Our data showthat the structure of the C-terminal region (NLS-TET-CTD) of full-length p53 tetramer is more compact than perceived by previous models. Moreover, even the DNA-free form is more compact and conformationally unchanged upon specific binding of a 26-mer RE-DNA. Our data most likely reflect the recognition mode (R-mode) of p53-DNA binding agreeing with previous data (25, 46). We are planning to apply our workflow to a variety of different DNA constructs varying in length as well as p53 specificity and affinity to contribute to the understanding of the underlying mechanisms for the transcriptional regulation of p53 by also capturing the sliding-mode (S-mode). The use of photo-reactive cross-linkers as well as HDX-MS will aid in resolving the complex dynamics upon p53 binding to different DNA molecules in addition to the already established distance restraints and labeling approach.

## Supporting information

Supplementary Inormation

## Acknowledgements

AS acknowledges financial support by the DFG (RTG 2467, project number 391498659 “Intrinsically Disordered Proteins – Molecular Principles, Cellular Functions, and Diseases”, INST 271/404-1 FUGG, INST 271/405-1 FUGG, and CRC 1423, (project number 421152132), the Federal Ministry for Economic Affairs and Energy (BMWi, ZIM project KK5096401SK0), the region of Saxony-Anhalt, and the Martin Luther University Halle-Wittenberg (Center for Structural Mass Spectrometry).

## Author Contributions

ADI and MK performed experiments, CT performed computational modeling, MK, CHI, CI, and JK analyzed data, CA, PLK, and AS supervised the study, CT, CA, and AS wrote the manuscript with input from all authors.

## Data Availability

All materials described in the manuscript, including all relevant raw data, will be freely available to any researcher wishing to use them for non-commercial purposes, without breaching participant confidentiality. All MS data are available via ProteomeXchange with identifier PXD037030.

## Supplementary Information

The Supplementary Information (.docx file) contains Figures S1 to S37 and Tables S3 and S4. Tables S1 and S2 are provided as .csv files (TableS1.csv and TableS2.csv).

