## Supplementary Inormation for "Structural Assessment of the Full-Length Wild-Type Tumor Suppressor Protein p53 by Mass Spectrometry-Guided Computational Modeling"

**Table of Contents**


| **Supplementary Figures** | page |
|---|---|
| Figure S1: Published tetrameric p53 models | 3 |
| Figure S2: SDS-PAGE analysis of quantitative XL-MS experiments | 4 |
| Figures S3-S16: Fragment ion mass spectra of interpeptide p53 cross-links | 5 |
| Figures S17-S31: Fragment ion mass spectra of p53 "dead-end" cross-links | 19 |
| Figure S32: AlphaFold2 prediction of tetrameric p53 | 34 |
| Figure S33: Refinement strategies | 35 |
| Figure S34: Scoring of XL-MS-derived distance restraints | 36 |
| Figure S35: Scoring of labeling efficiency | 37 |
| Figure S36: XL-MS and footprinting data of p53 tetramer | 38 |
| Figure S37: XL-MS and footprinting data mapped in existing p53 models | 39 |

| **Supplementary Tables** | |
|---|---|
| **Table S1:** $BS^2G$ and DSBU cross-linking data (file TableS1.csv) | 40 |
| **Table S2:** SASA scoring of labeled lysine residues (file TableS2.csv) | 40 |
| **Table S3:** $BS^2G$ cross-links of p53 (+/- RE-DNA) | 40 |
| **Table S4:** Footprinting of p53 (+/- RE-DNA) | 41 |

### Supplementary Figures

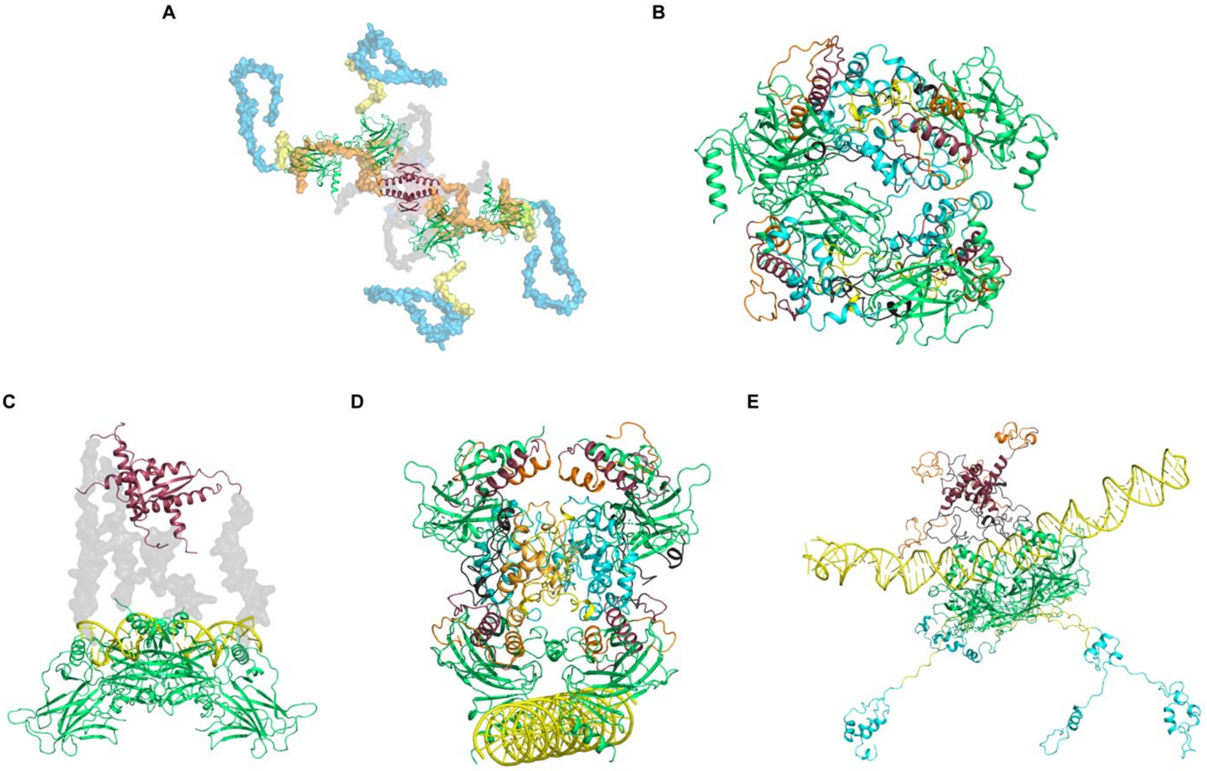


**Figure S1:** Tetrameric p53 models in cartoon presentation. DNA-free p53: (A) Tidow *et al*, 2007 (20), (B) Okorokov *et al*, 2006 (19). DNA-bound p53: (C) Tidow *et al*, 2007 (20), (D) Okorokov *et al*, 2006 (19), and (E) Demir *et al*, 2017 (25). TAD is presented in light blue, PRR is yellow, DBD in green, NLS in orange, TET in red, and CTD in dark grey.

**A**

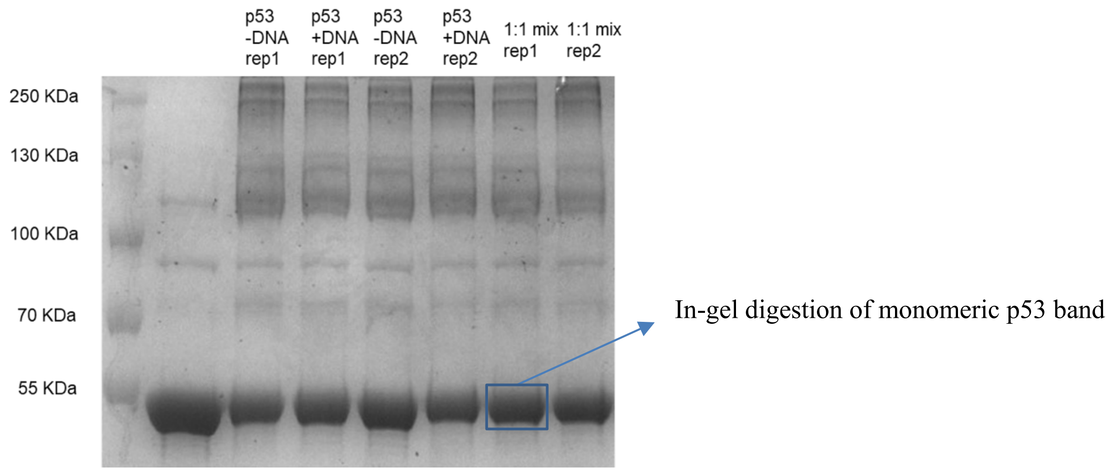


**B**

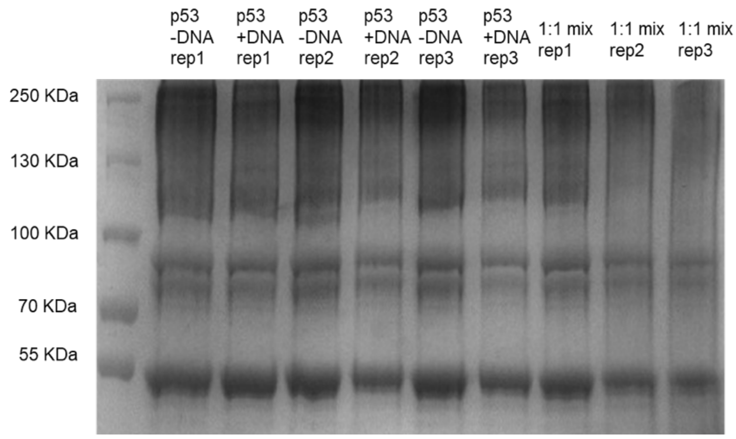


**Figures S2:** SDS-PAGE analysis of quantitative XL-MS experiments of p53 with BS$^2$G-D$_0$/D$_4$. (A) p53 without DNA: BS$^2$G-D$_0$, with DNA: BS$^2$G-D$_4$. (B) p53 without DNA: BS$^2$G-D$_4$, with DNA: BS$^2$G-D$_0$.

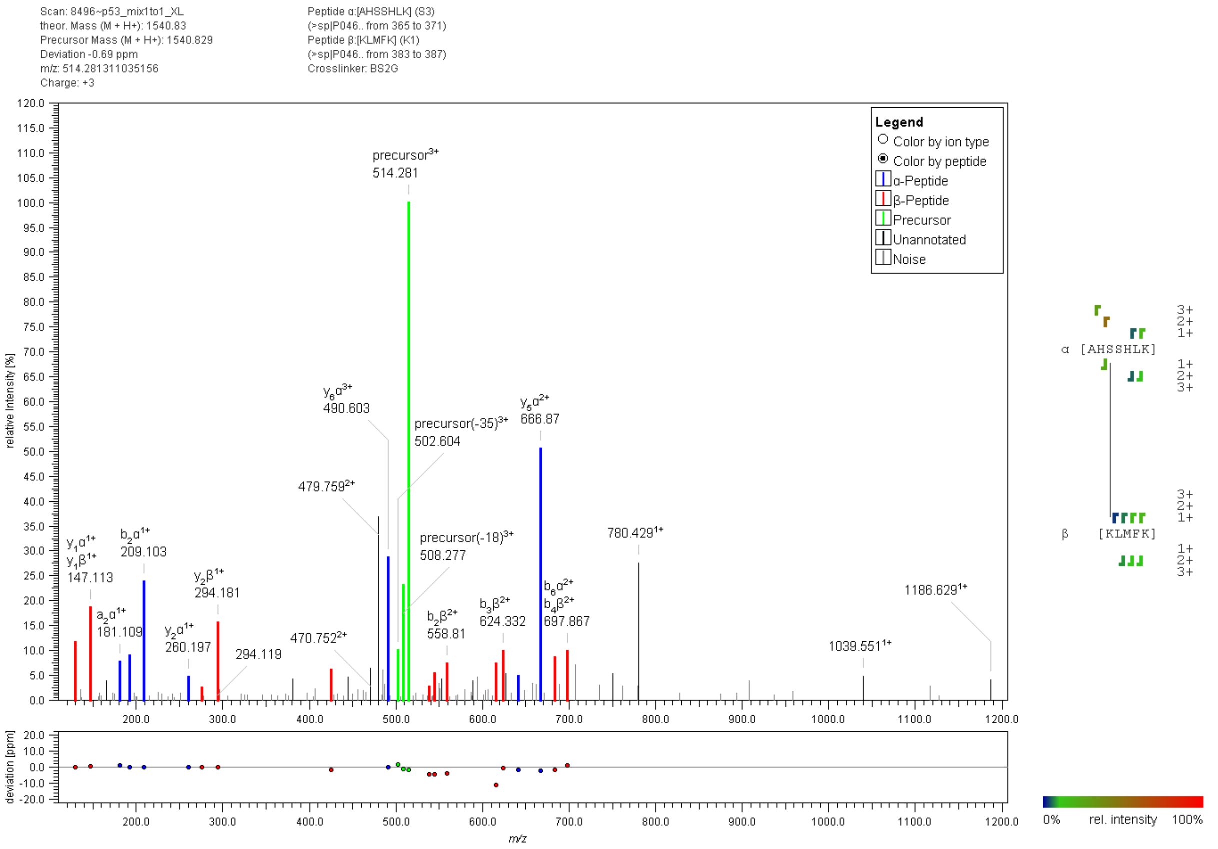


**Figure S3:** Fragment ion mass spectrum of an interpeptide p53 cross-link with $BS^2G$. Signal assignment was performed with MeroX 2.0.1.7 (39).

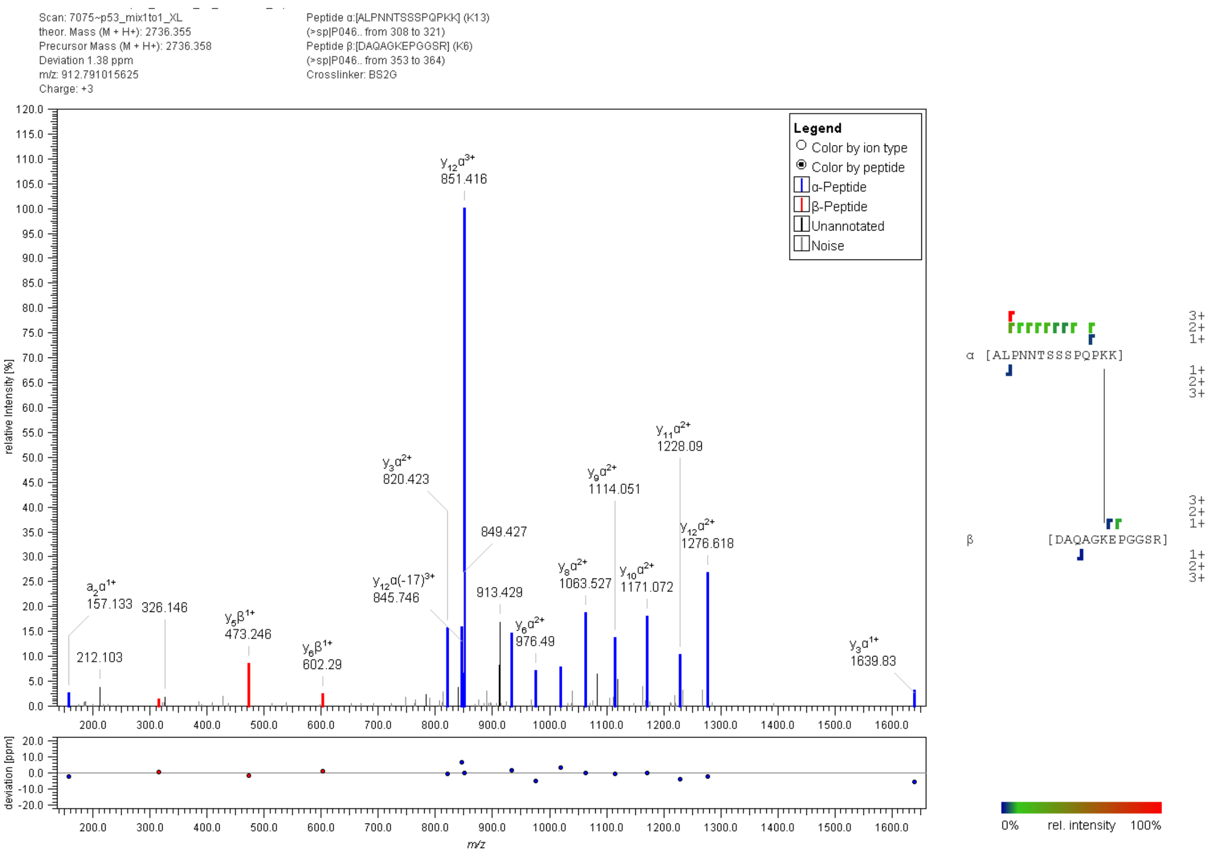


**Figure S4:** Fragment ion mass spectrum of an interpeptide p53 cross-link with $BS^2G$. Signal assignment was performed with MeroX 2.0.1.7 (39).

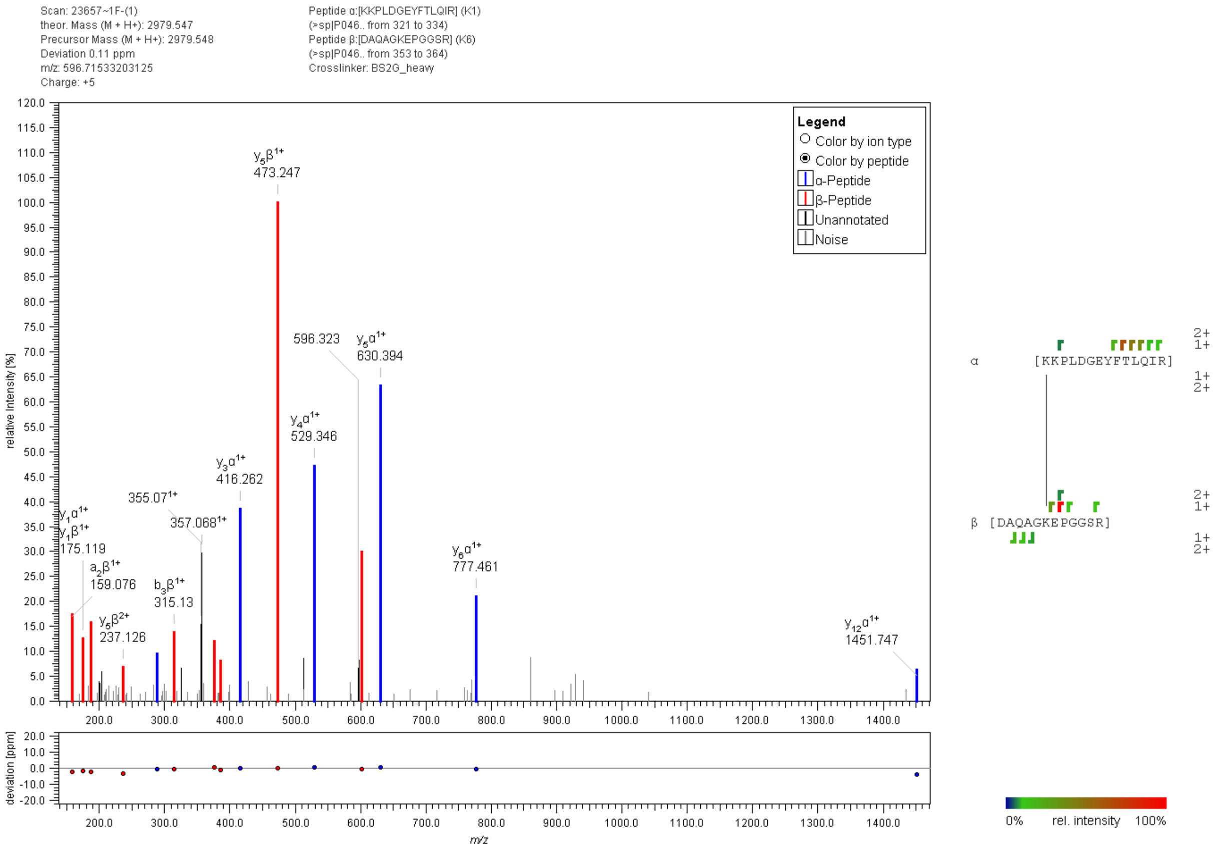


**Figure S5:** Fragment ion mass spectrum of an interpeptide p53 cross-link with $BS^2G$. Signal assignment was performed with MeroX 2.0.1.7 (39).

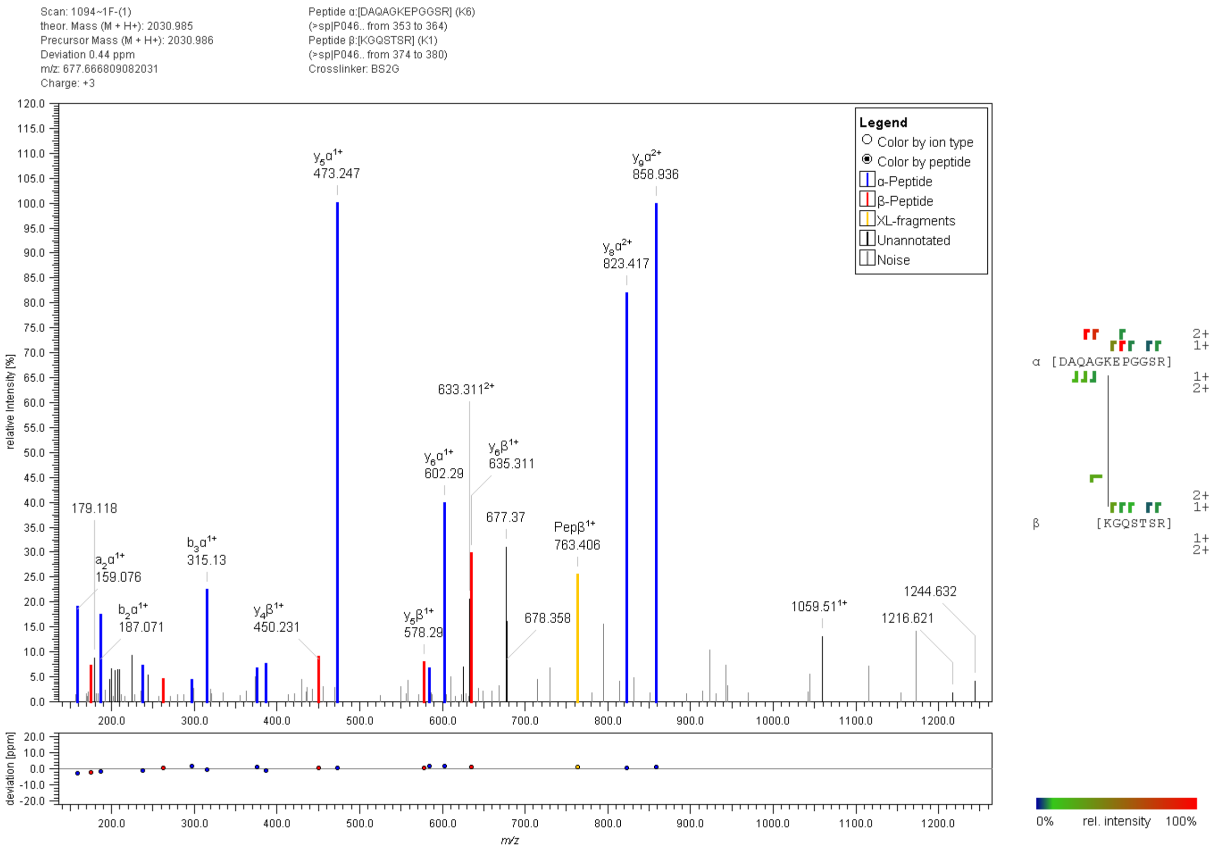


**Figure S6:** Fragment ion mass spectrum of an interpeptide p53 cross-link with BS$^2$G. Signal assignment was performed with MeroX 2.0.1.7 (39).

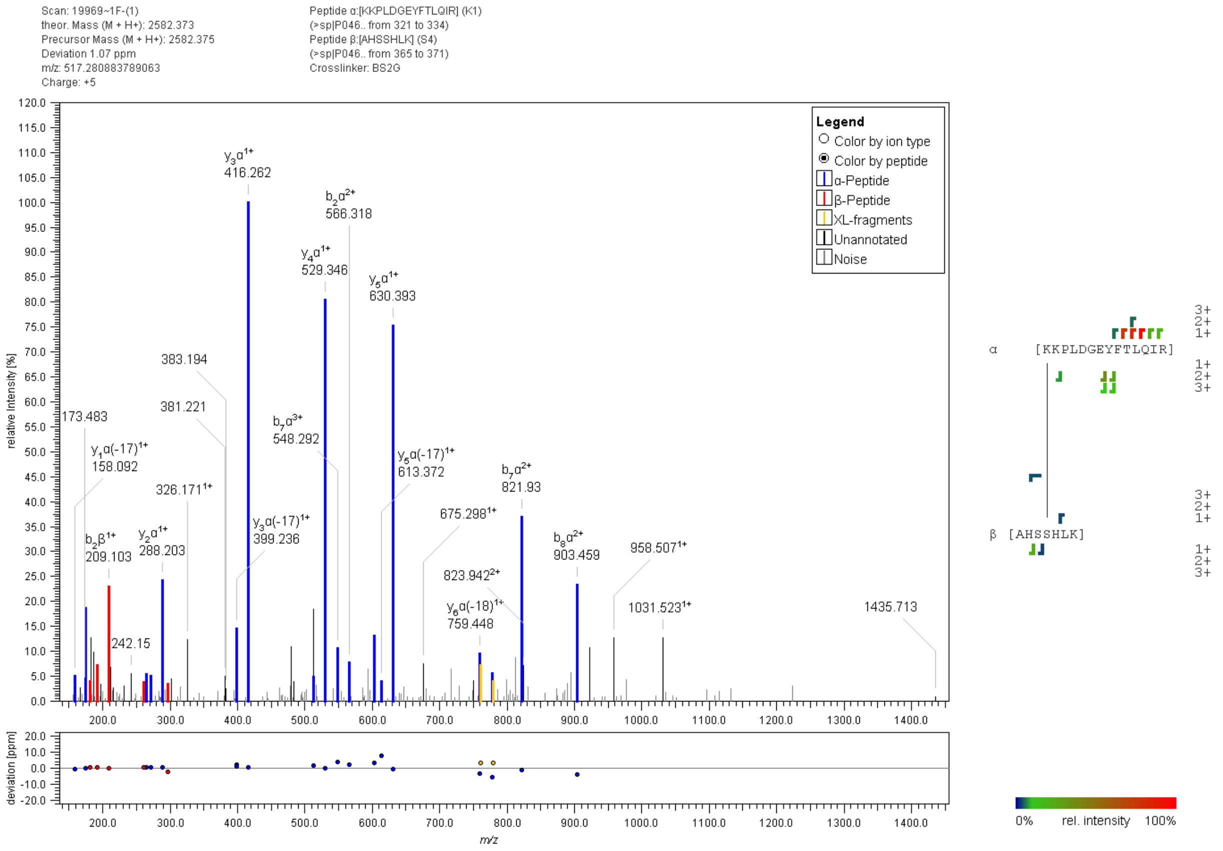


**Figure S7:** Fragment ion mass spectrum of an interpeptide p53 cross-link with BS$^2$G. Signal assignment was performed with MeroX 2.0.1.7 (39).

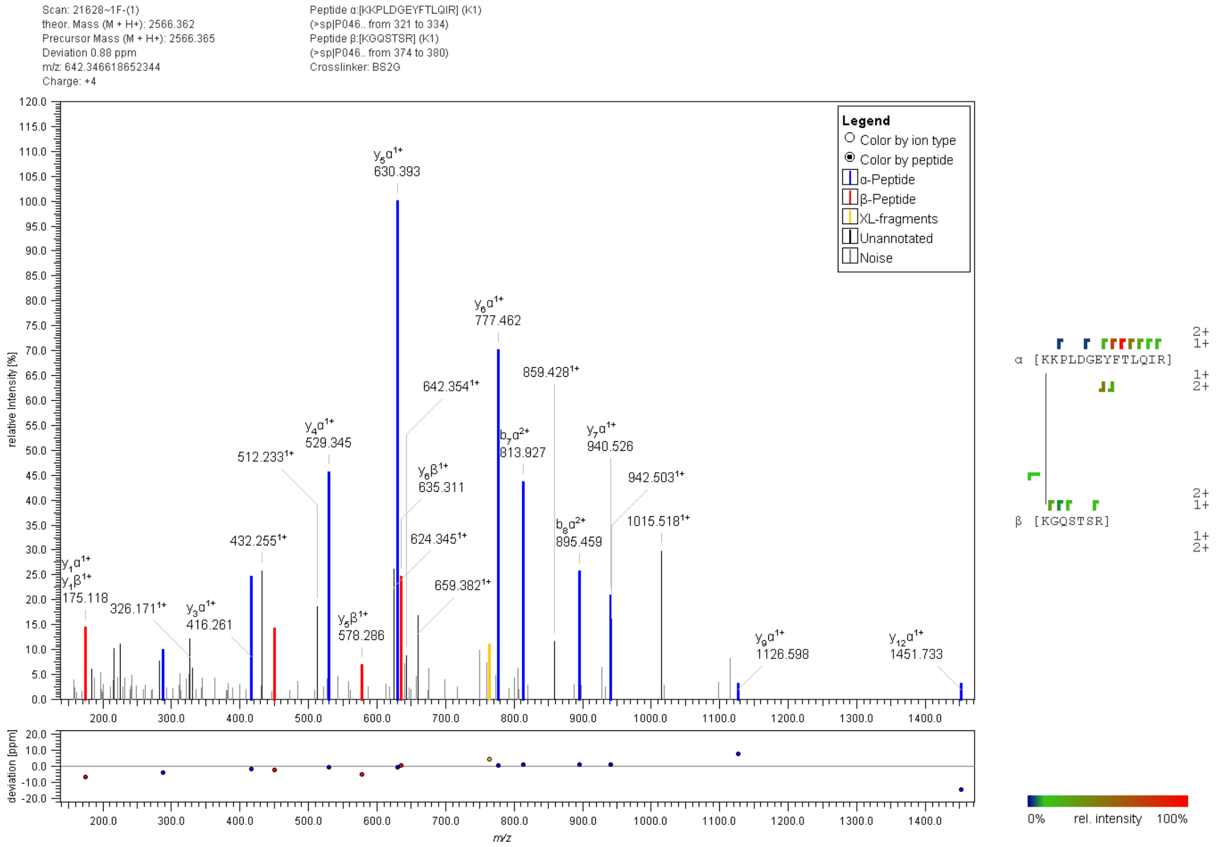


**Figure S8:** Fragment ion mass spectrum of an interpeptide p53 cross-link with BS$^2$G. Signal assignment was performed with MeroX 2.0.1.7 (39).

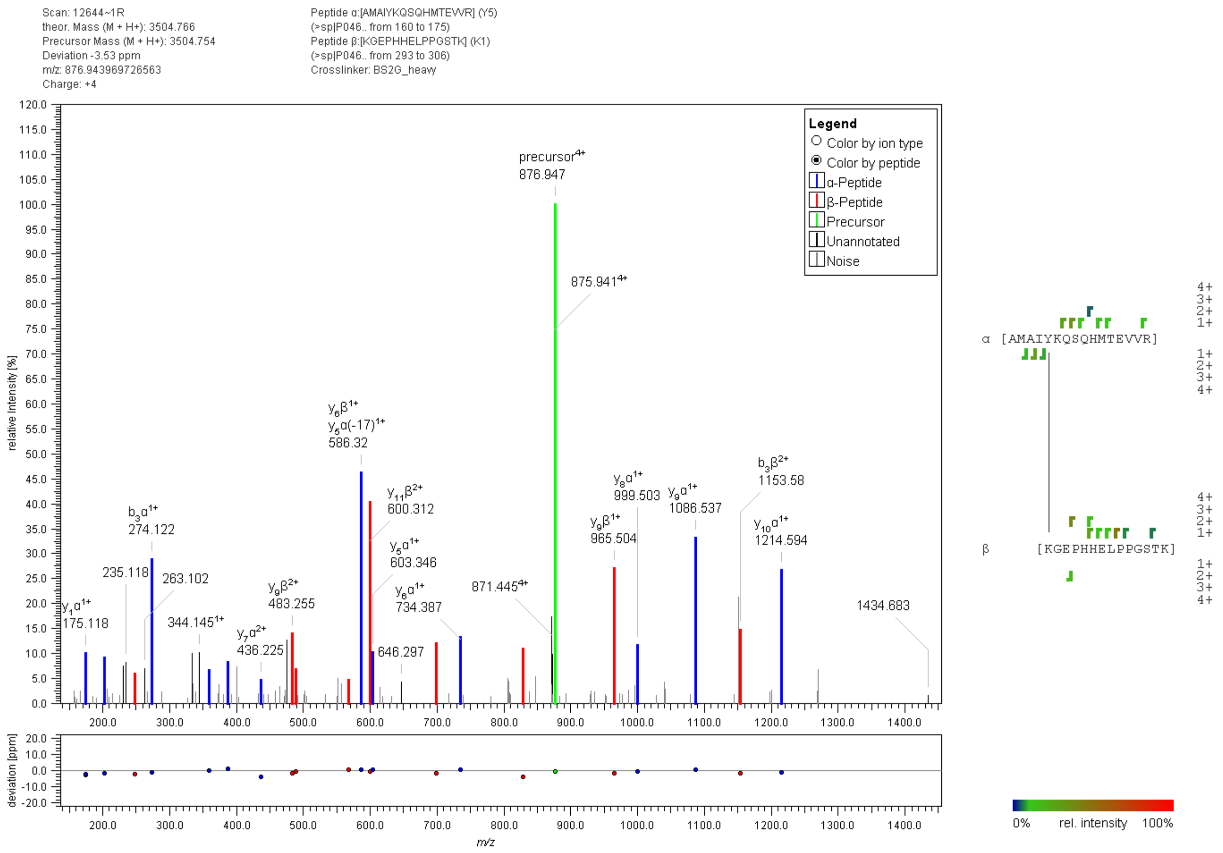


**Figure S9:** Fragment ion mass spectrum of an interpeptide p53 cross-link with BS$^2$G. Signal assignment was performed with MeroX 2.0.1.7 (39).

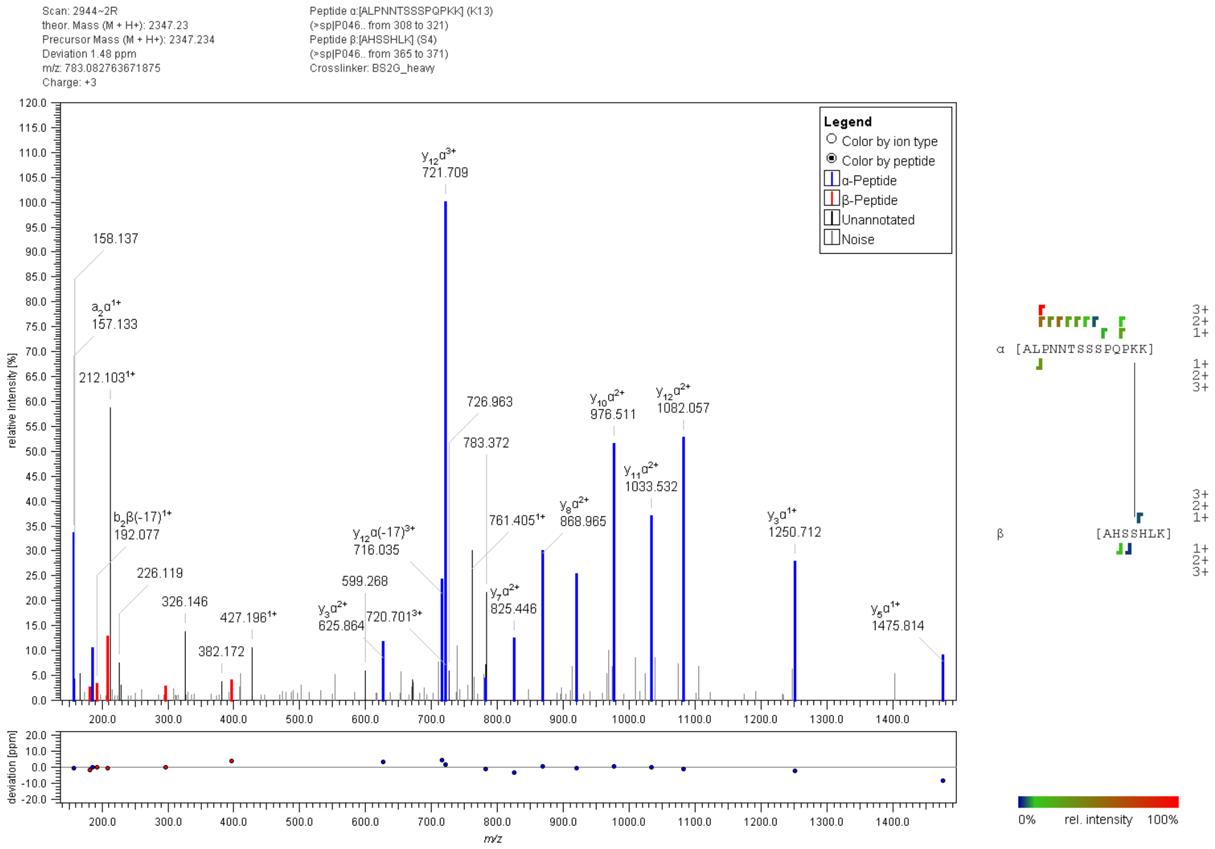


**Figure S10:** Fragment ion mass spectrum of an interpeptide p53 cross-link with BS$^2$G. Signal assignment was performed with MeroX 2.0.1.7 (39).

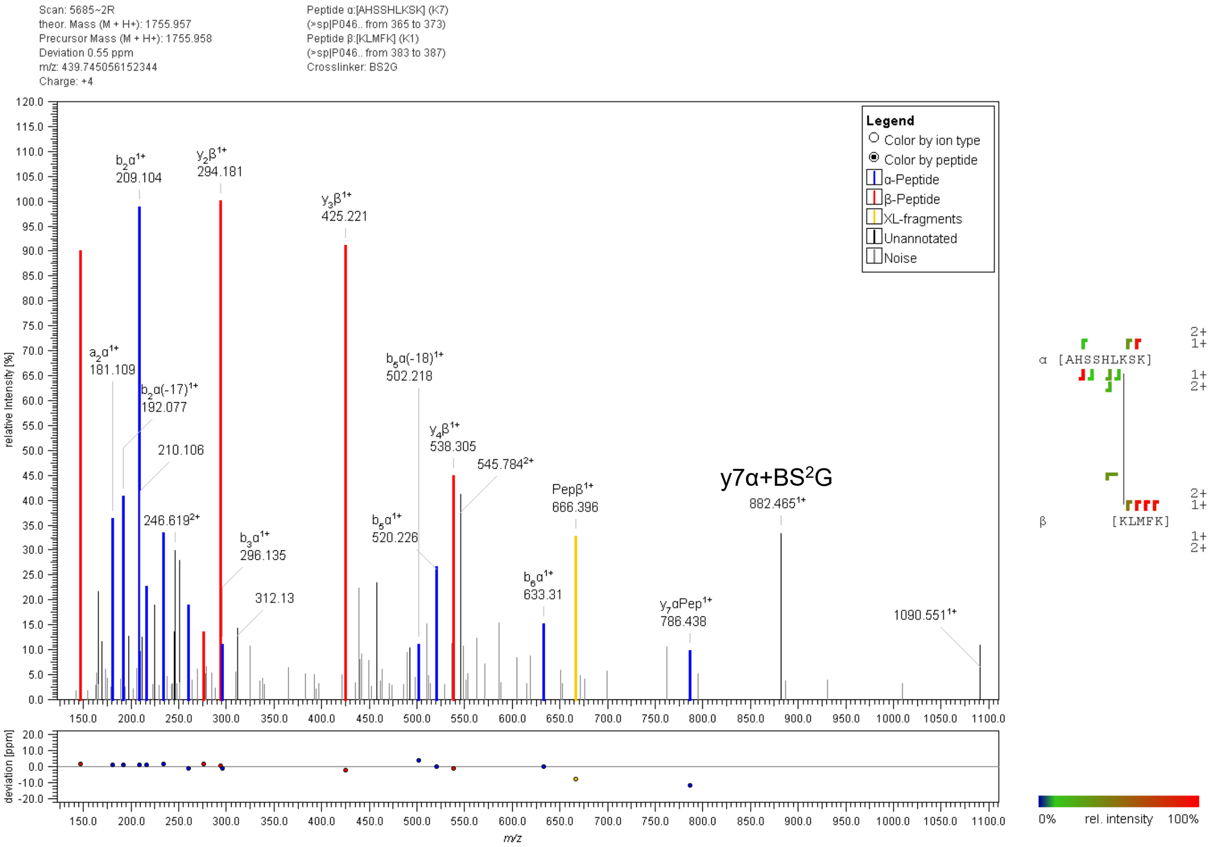


**Figure S11:** Fragment ion mass spectrum of an interpeptide p53 cross-link with BS$^2$G. Signal assignment was performed with MeroX 2.0.1.7 (39).

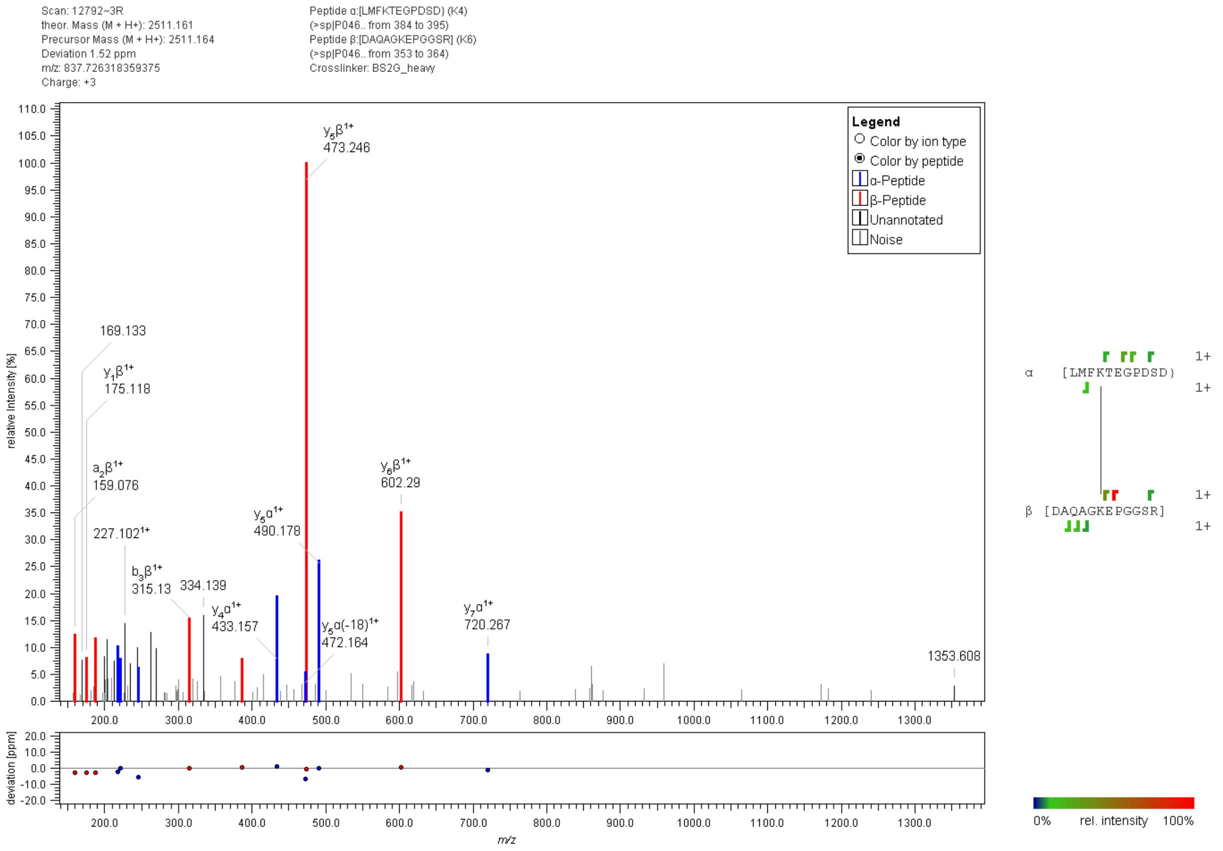


**Figure S12:** Fragment ion mass spectrum of an interpeptide p53 cross-link with $BS^2G$. Signal assignment was performed with MeroX 2.0.1.7 (39).

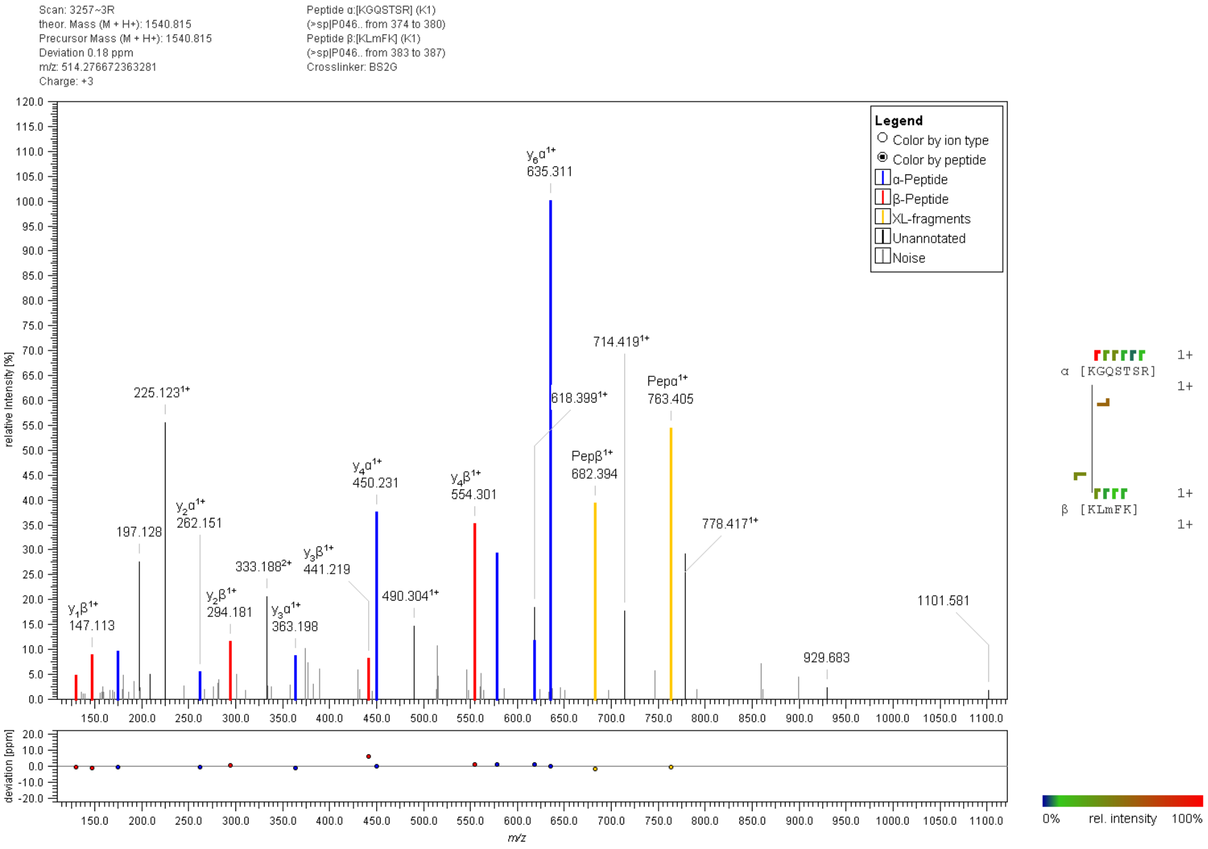


**Figure S13:** Fragment ion mass spectrum of an interpeptide p53 cross-link with BS$^2$G. Signal assignment was performed with MeroX 2.0.1.7 (39).

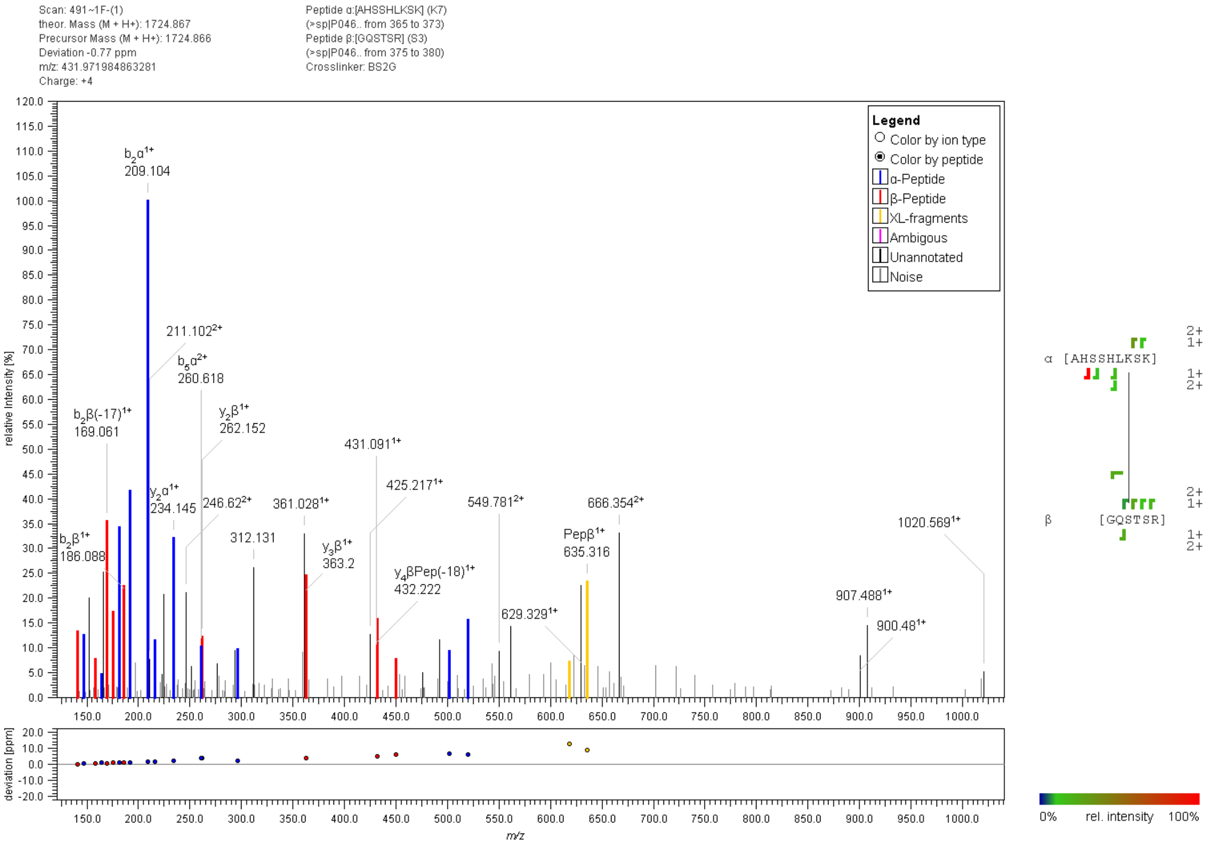


**Figure S14:** Fragment ion mass spectrum of an interpeptide p53 cross-link with BS$^2$G. Signal assignment was performed with MeroX 2.0.1.7 (39).

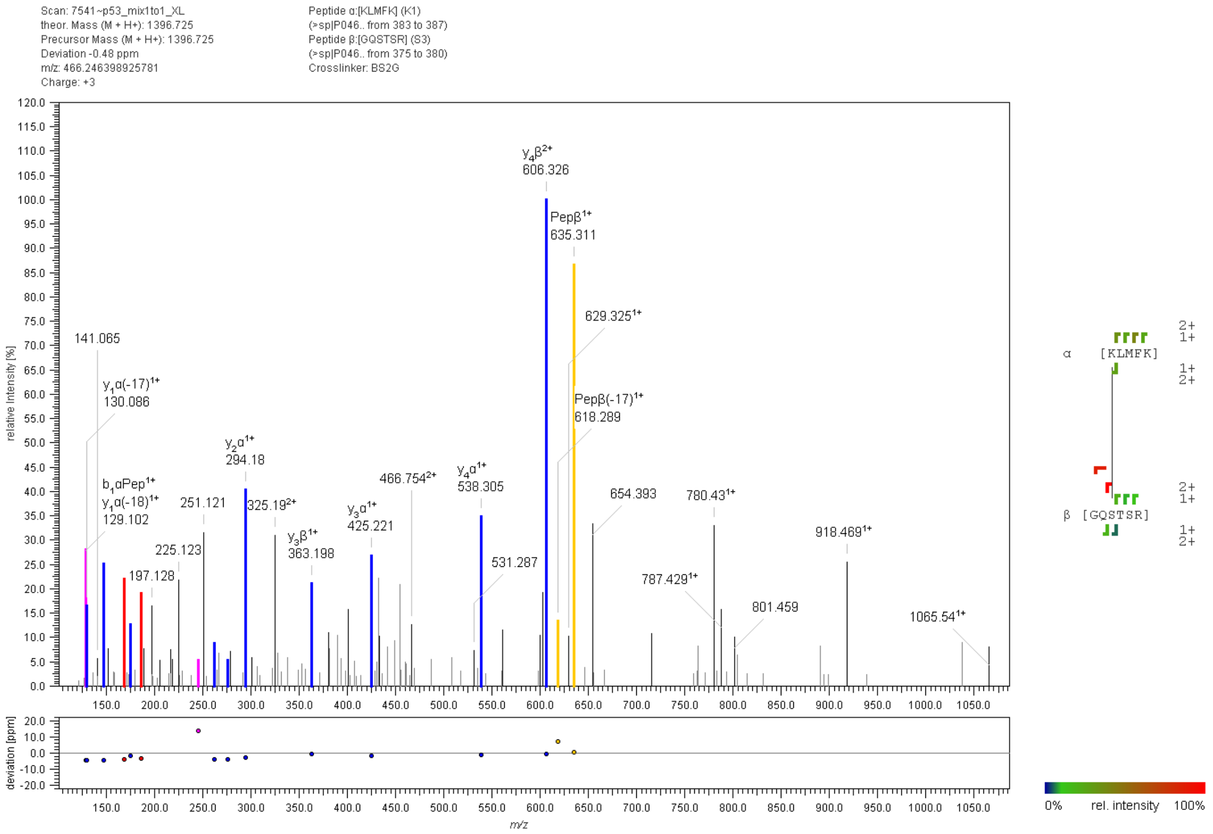


**Figure S15:** Fragment ion mass spectrum of an interpeptide p53 cross-link with BS$^2$G. Signal assignment was performed with MeroX 2.0.1.7 (39).

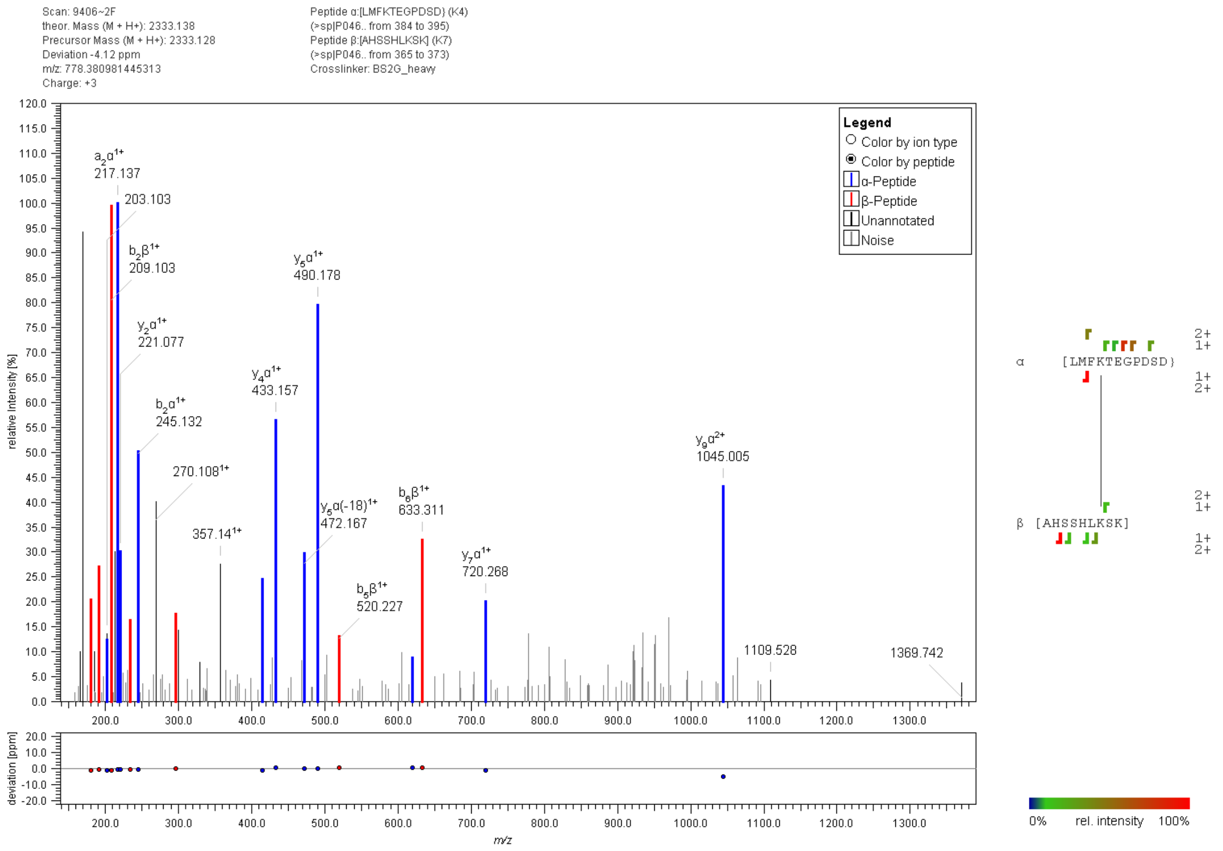


**Figure S16:** Fragment ion mass spectrum of an interpeptide p53 cross-link with BS$^2$G. Signal assignment was performed with MeroX 2.0.1.7 (39).

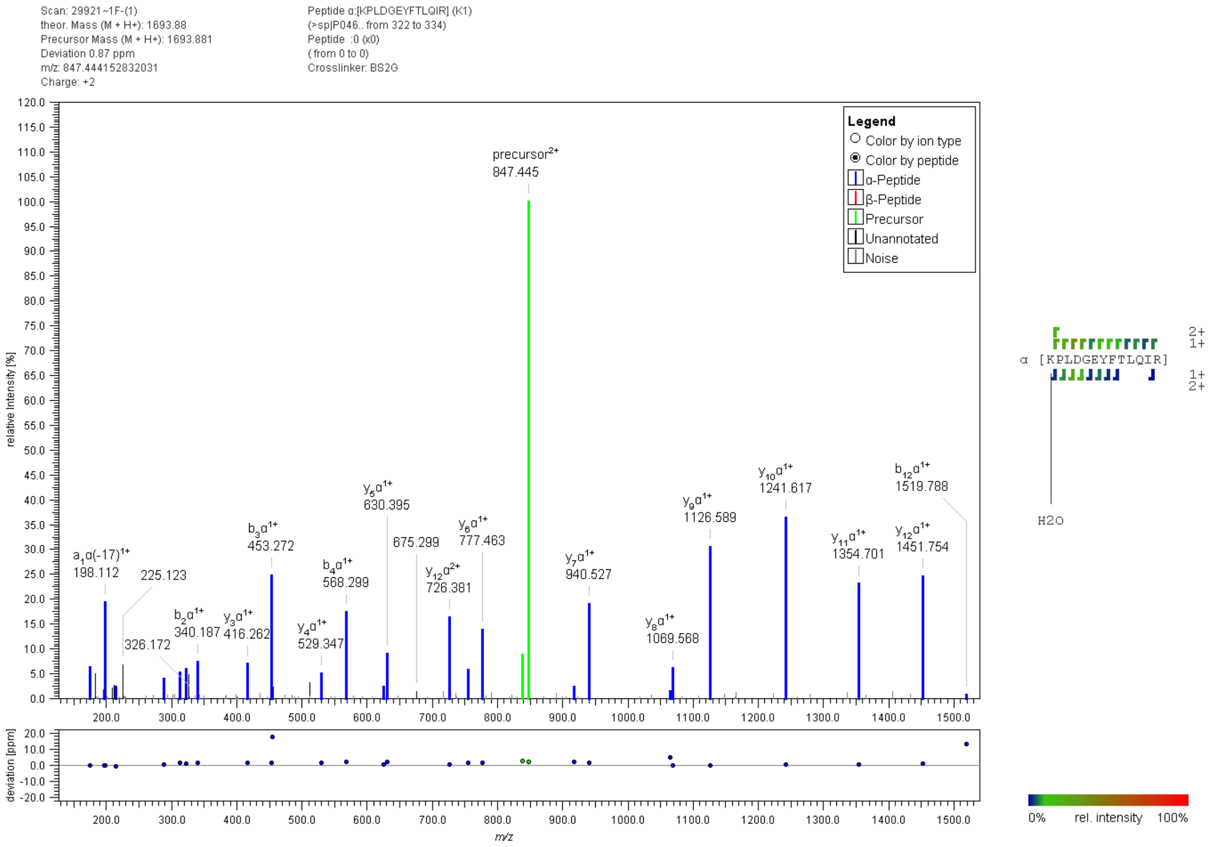


**Figure S17:** Fragment ion mass spectrum of a p53 "dead-end" cross-link with BS$^2$G. Signal assignment was performed with MeroX 2.0.1.7 (39).

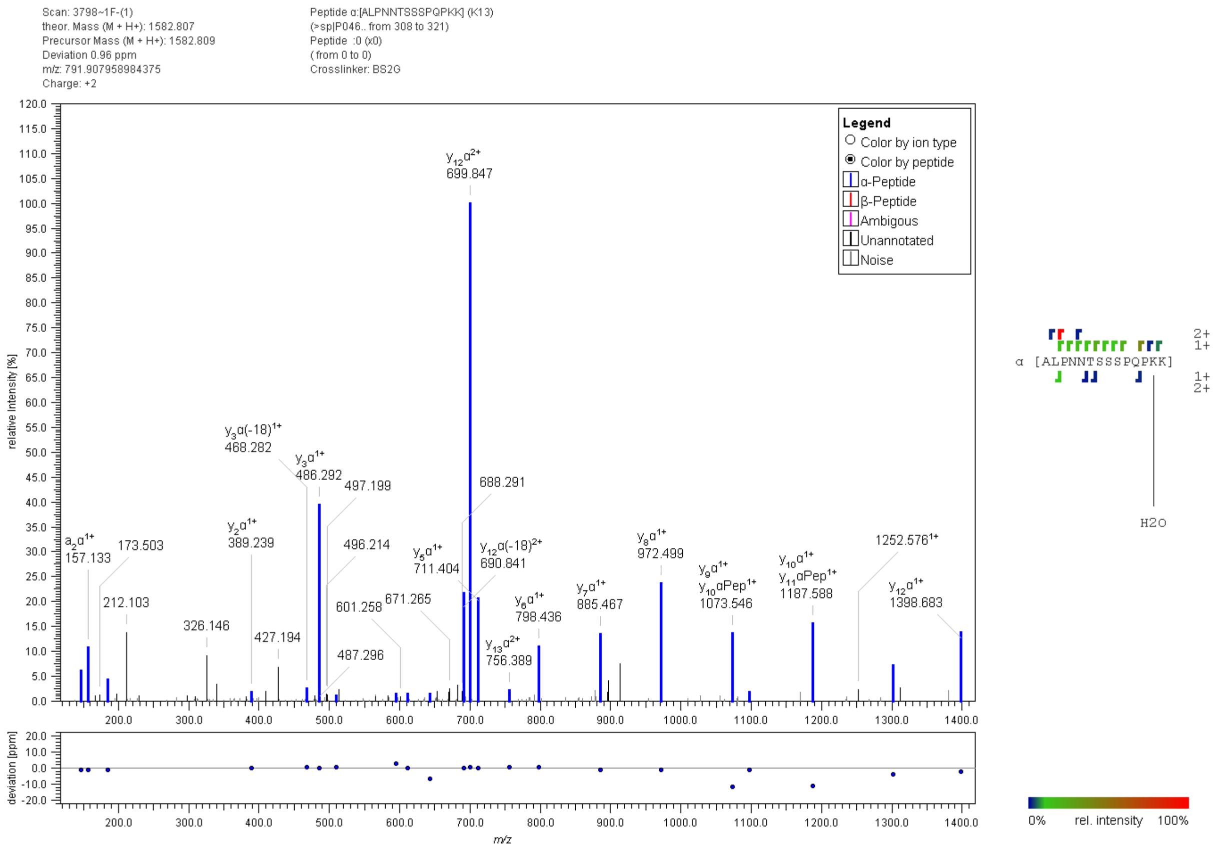


**Figure S18:** Fragment ion mass spectrum of a p53 "dead-end" cross-link with BS$^2$G. Signal assignment was performed with MeroX 2.0.1.7 (39).

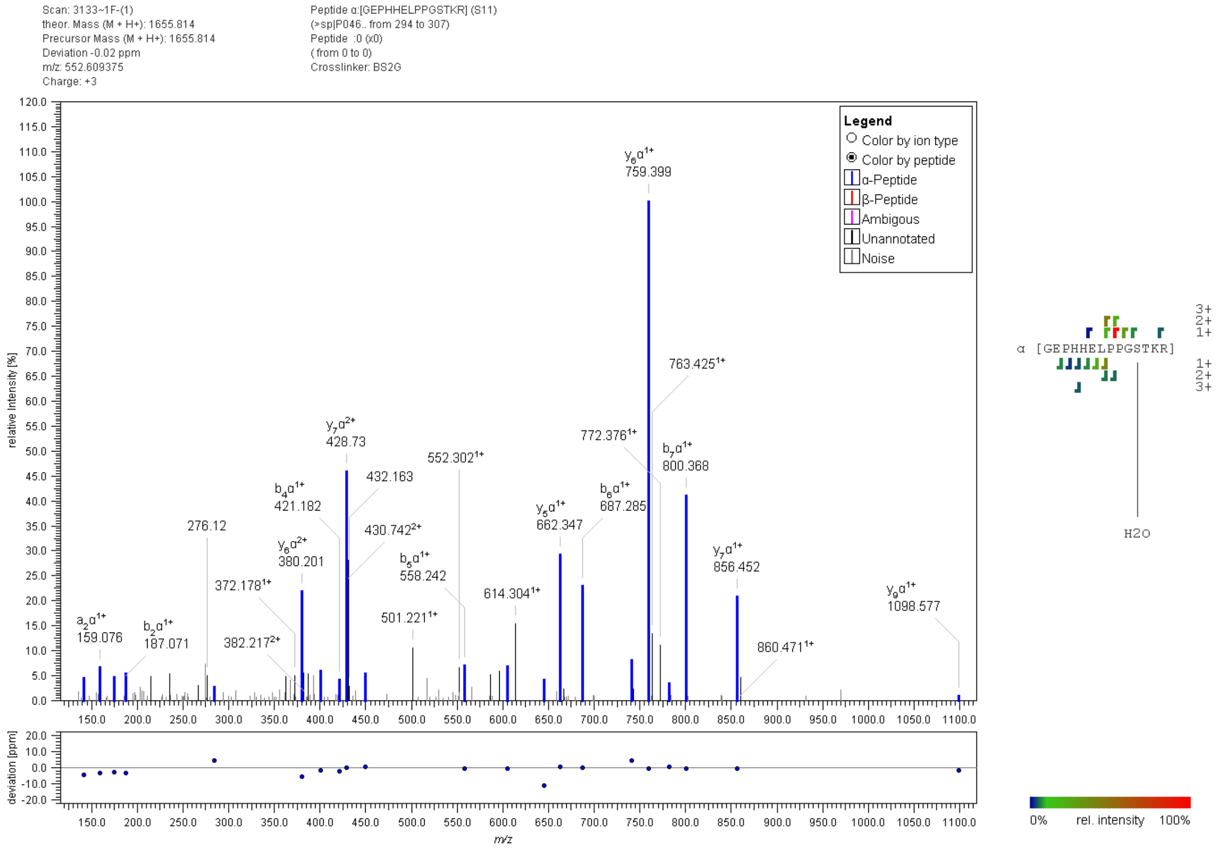


**Figure S19:** Fragment ion mass spectrum of a p53 "dead-end" cross-link with BS$^2$G. Signal assignment was performed with MeroX 2.0.1.7 (39).

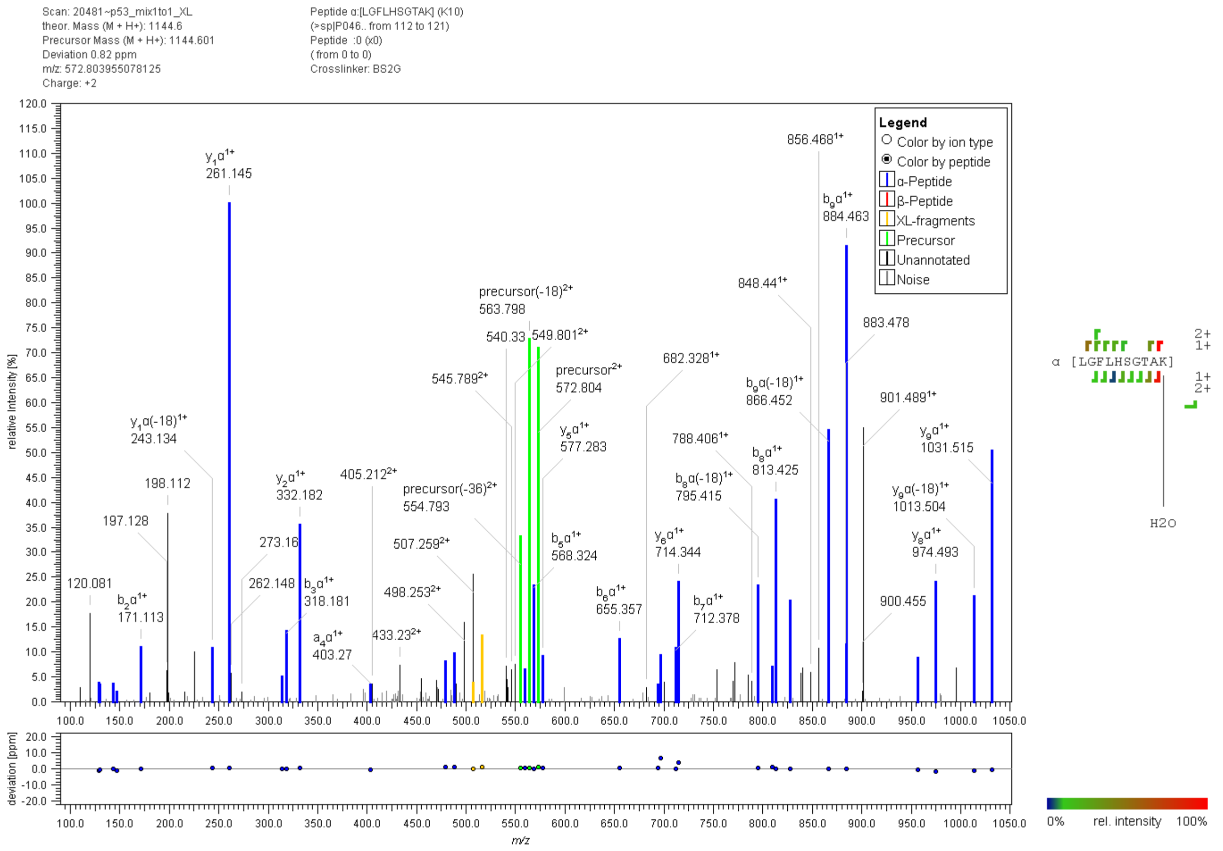


**Figure S20:** Fragment ion mass spectrum of a p53 "dead-end" cross-link with BS$^2$G. Signal assignment was performed with MeroX 2.0.1.7 (39).

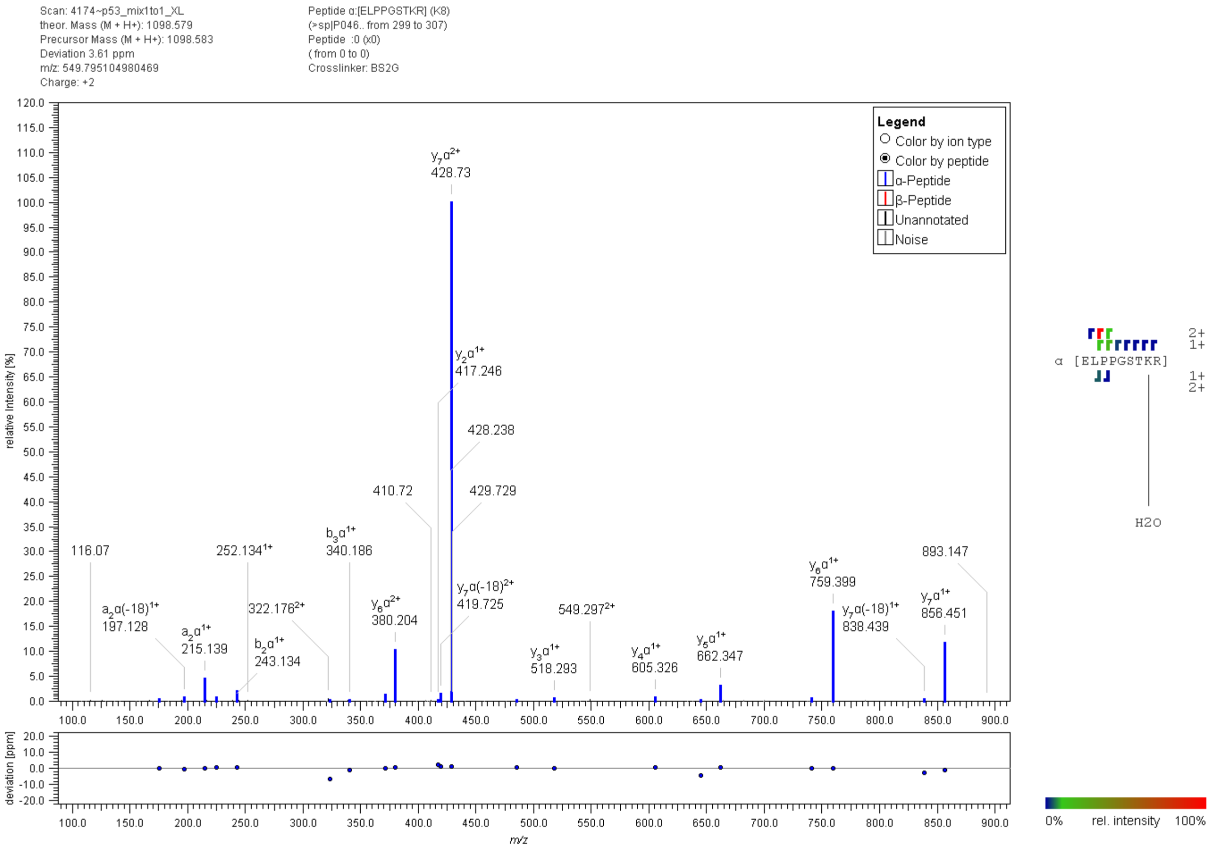


**Figure S21:** Fragment ion mass spectrum of a p53 "dead-end" cross-link with $BS^2G$. Signal assignment was performed with MeroX 2.0.1.7 (39).

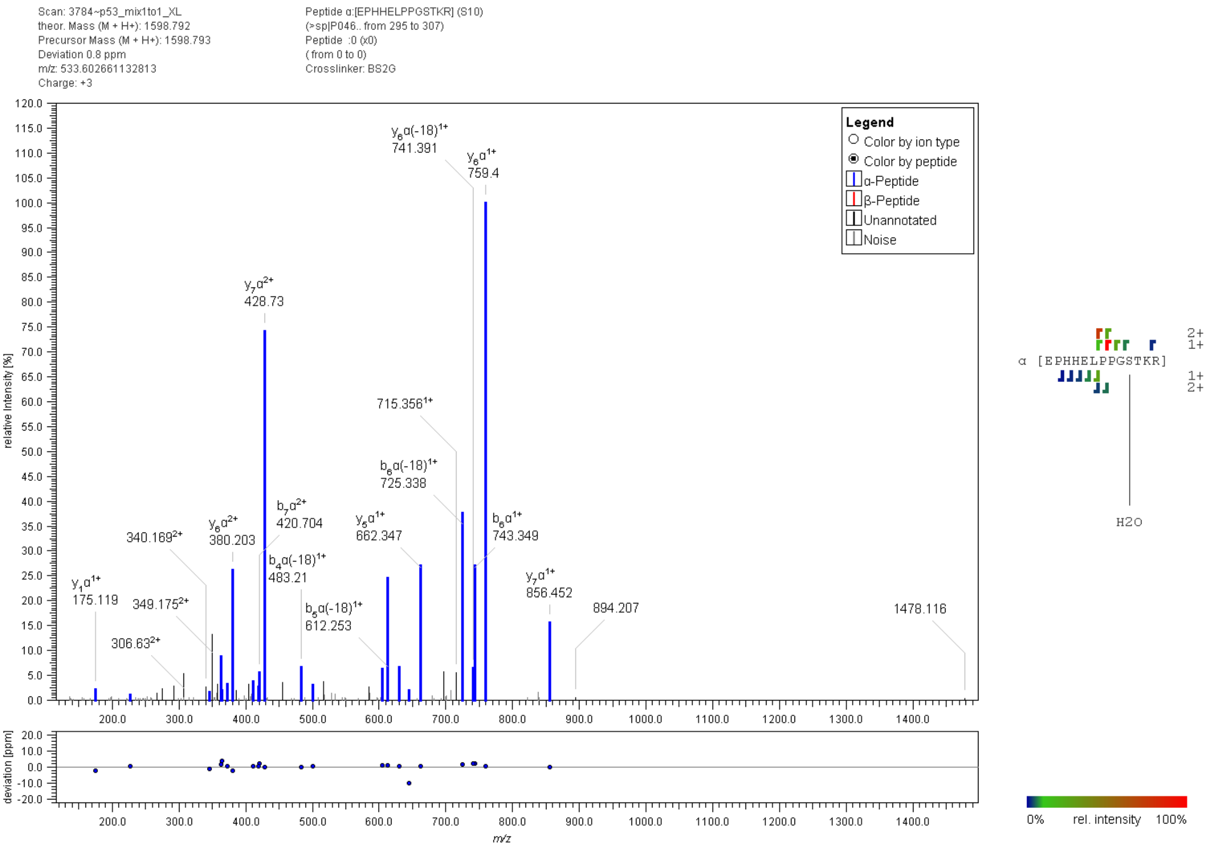


**Figure S22:** Fragment ion mass spectrum of a p53 "dead-end" cross-link with $BS^2G$. Signal assignment was performed with MeroX 2.0.1.7 (39).

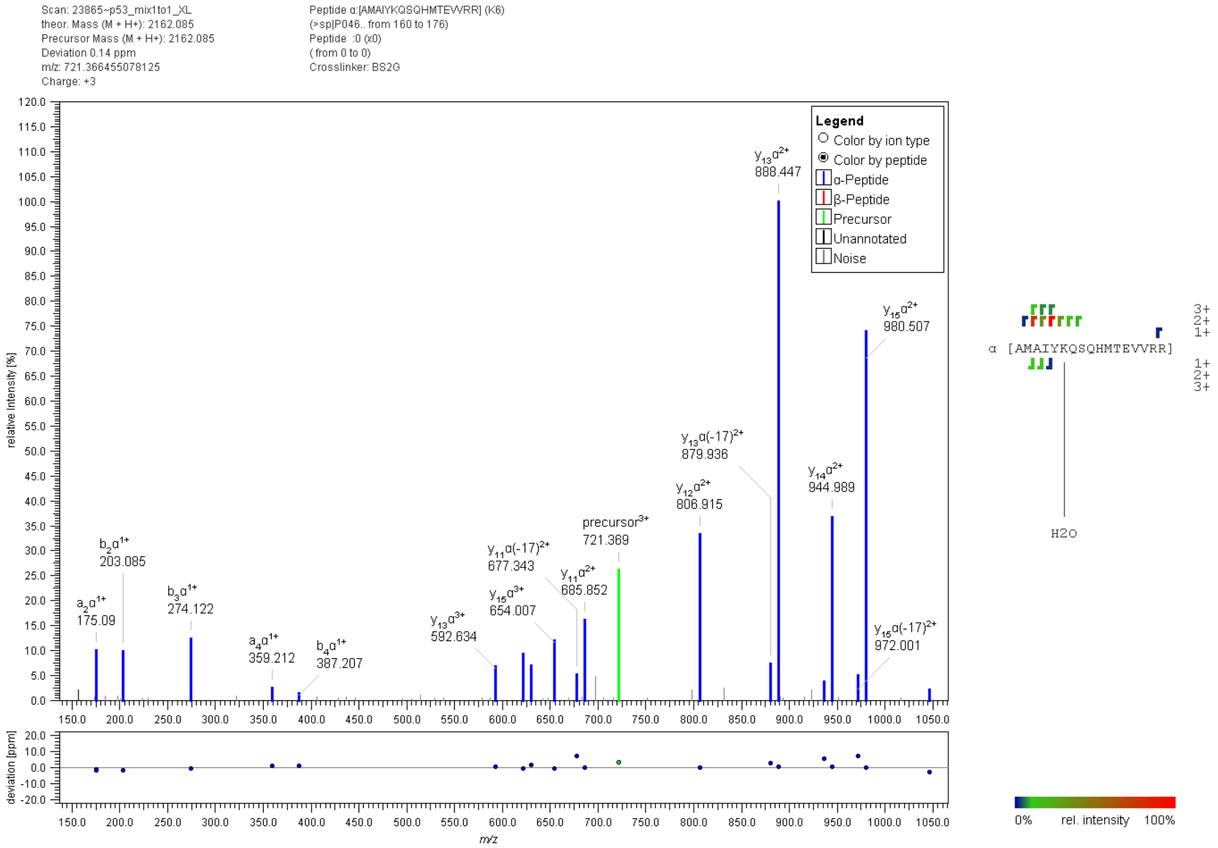


**Figure S23:** Fragment ion mass spectrum of a p53 "dead-end" cross-link with BS$^2$G. Signal assignment was performed with MeroX 2.0.1.7 (39).

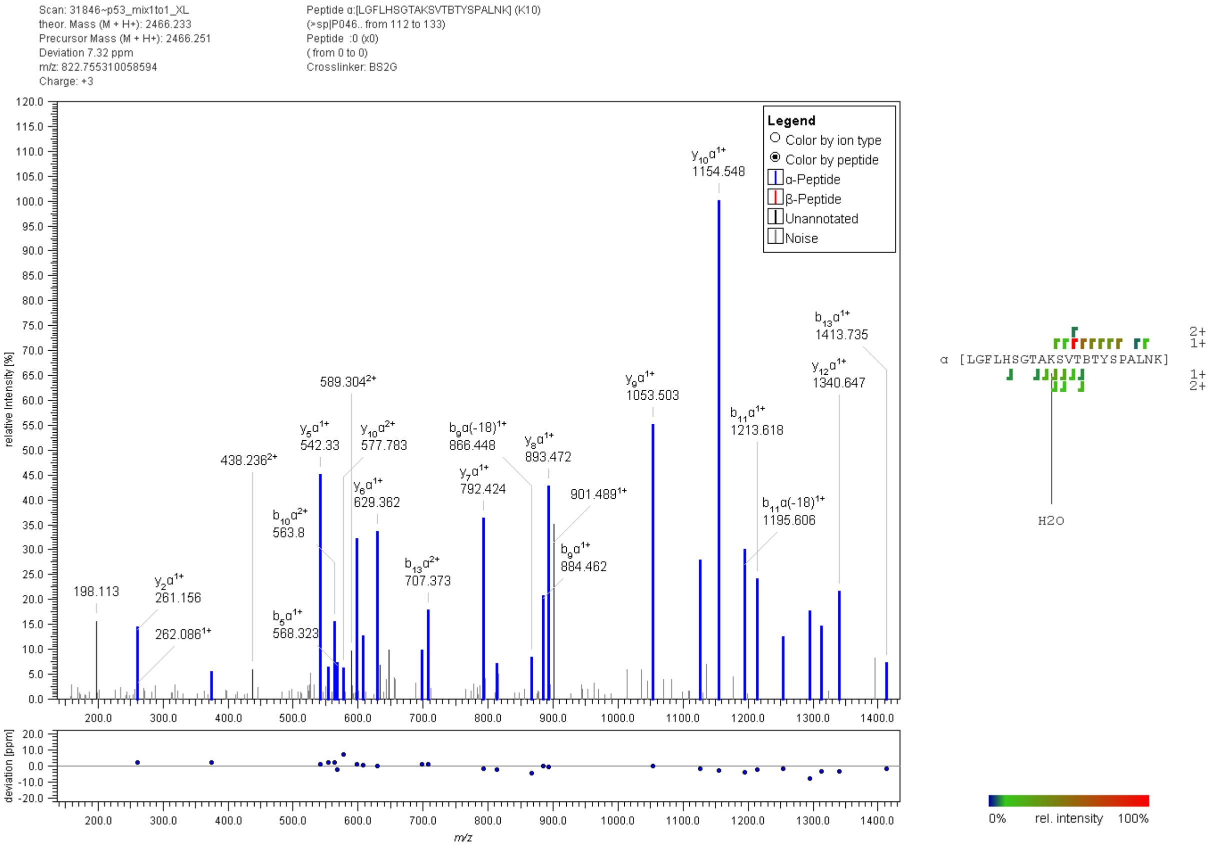


**Figure S24:** Fragment ion mass spectrum of a p53 “dead-end” cross-link with BS$^2$G. Signal assignment was performed with MeroX 2.0.1.7 (39).

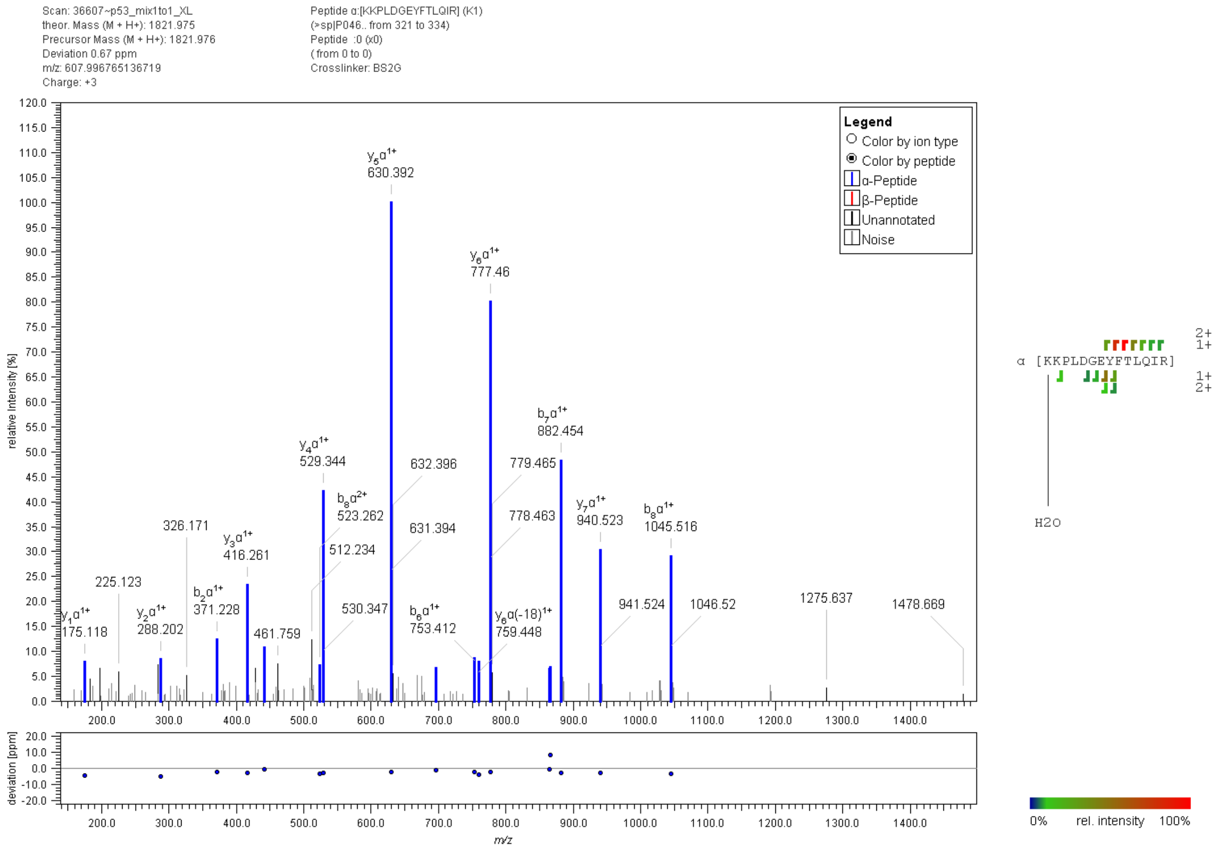


**Figure S25:** Fragment ion mass spectrum of a p53 “dead-end” cross-link with $BS^2G$. Signal assignment was performed with MeroX 2.0.1.7 (39).

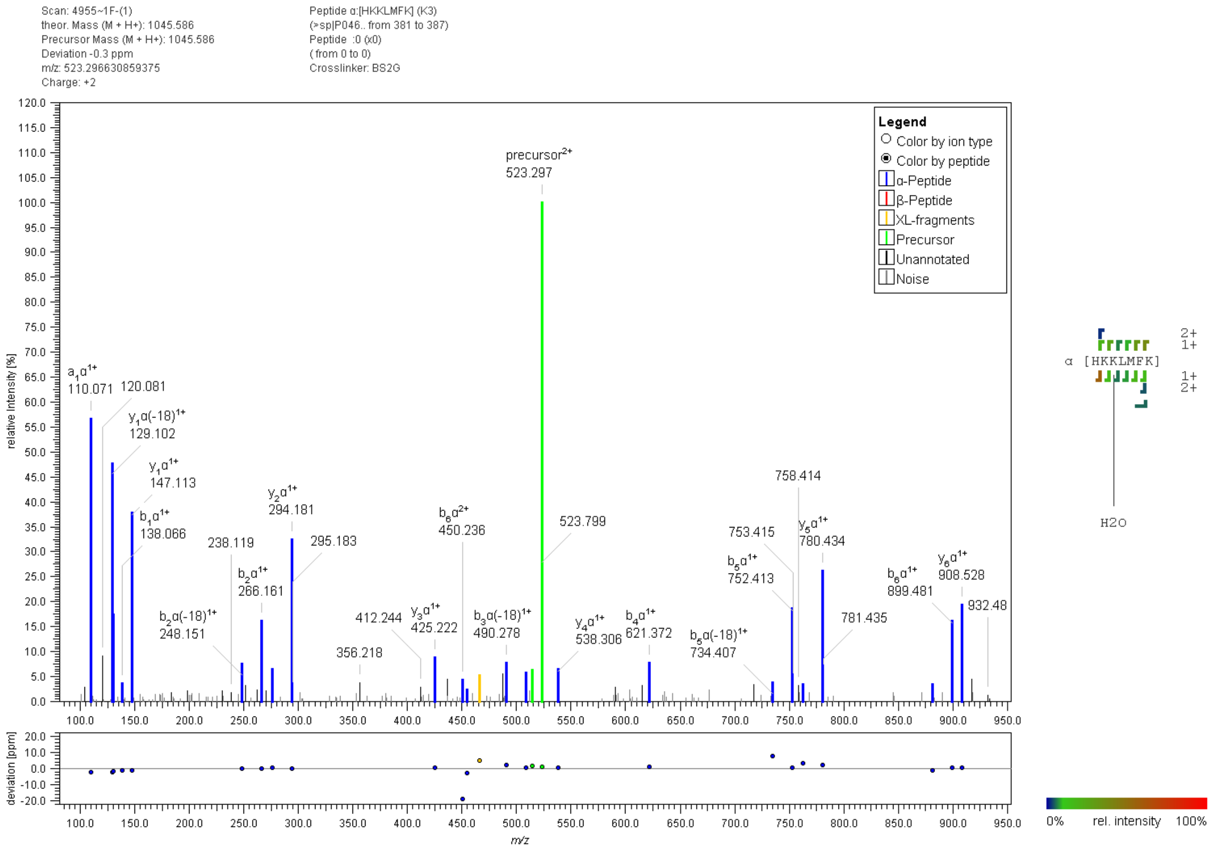


**Figure S26:** Fragment ion mass spectrum of a p53 "dead-end" cross-link with BS$^2$G. Signal assignment was performed with MeroX 2.0.1.7 (39).

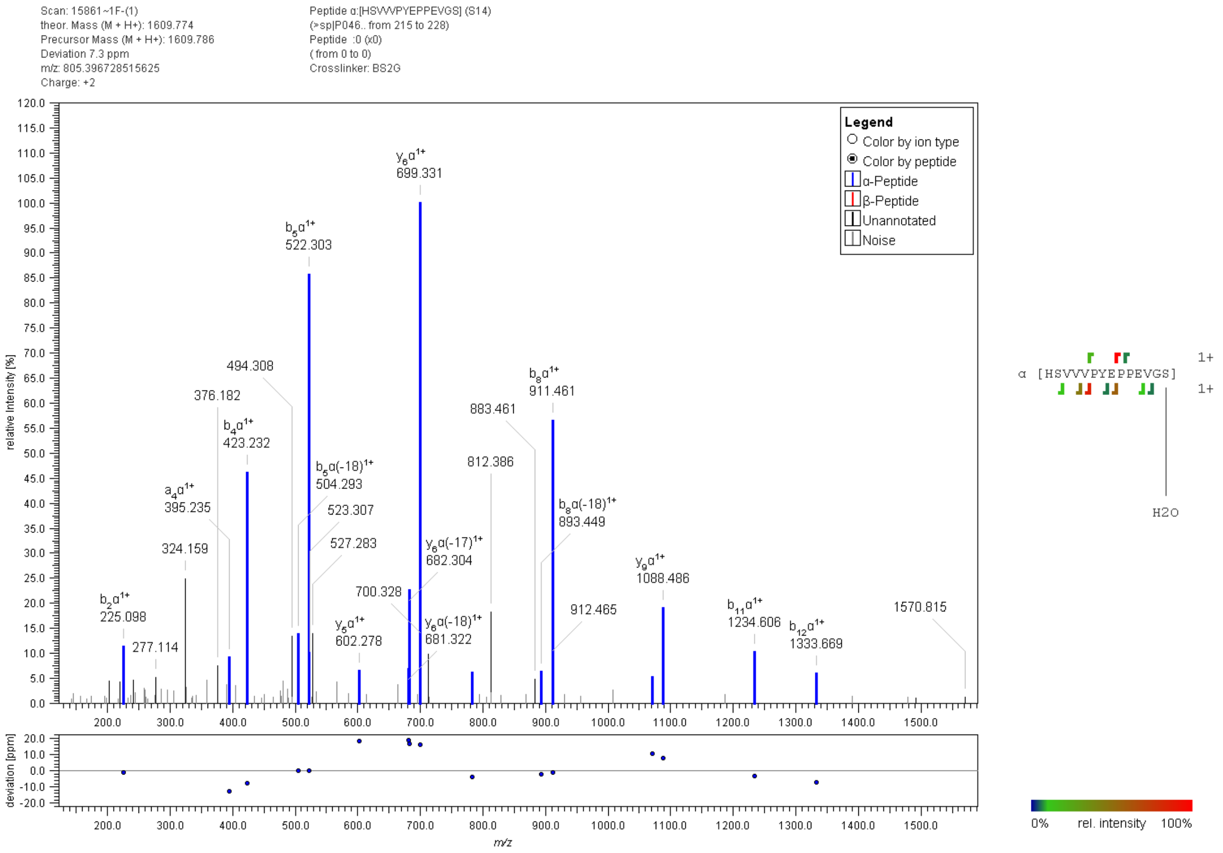


**Figure S27:** Fragment ion mass spectrum of a p53 “dead-end” cross-link with BS$^2$G. Signal assignment was performed with MeroX 2.0.1.7 (39).

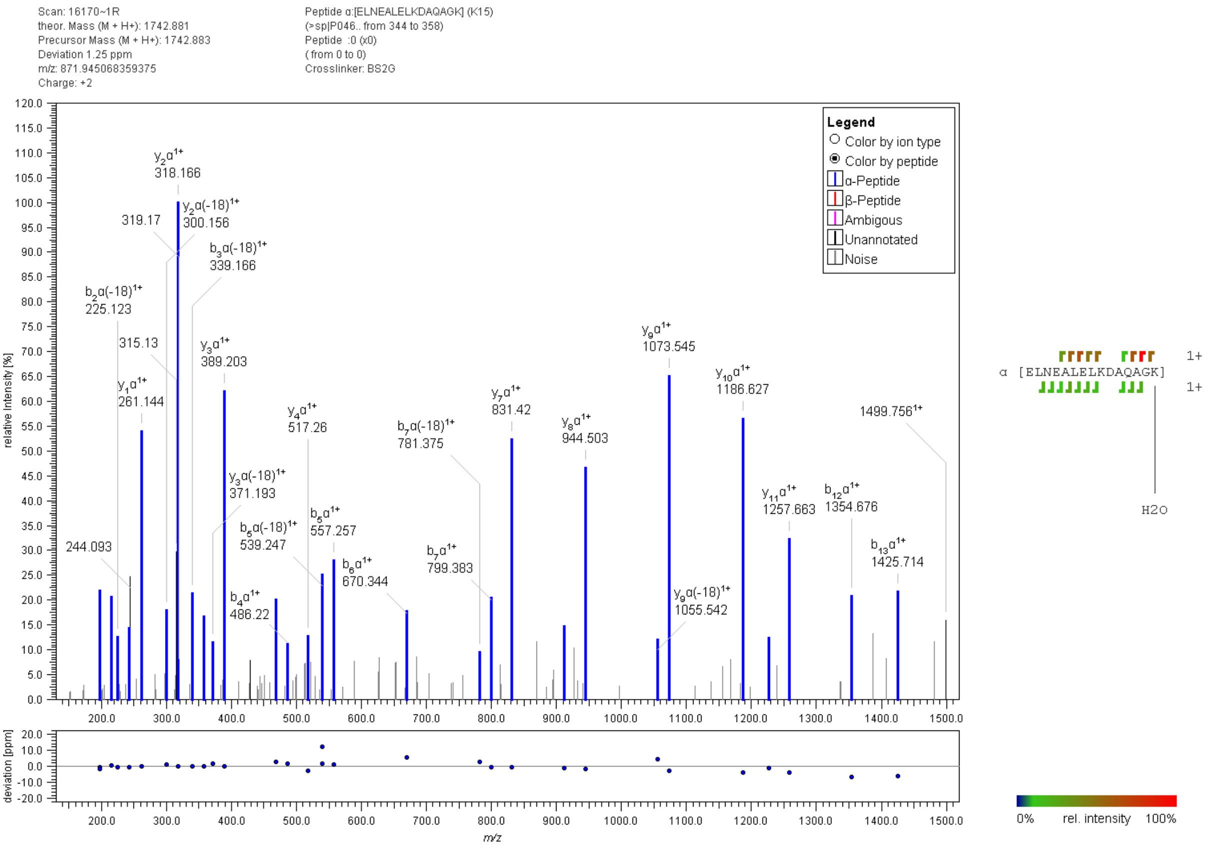


**Figure S28:** Fragment ion mass spectrum of a p53 “dead-end” cross-link with BS$^2$G. Signal assignment was performed with MeroX 2.0.1.7 (39).

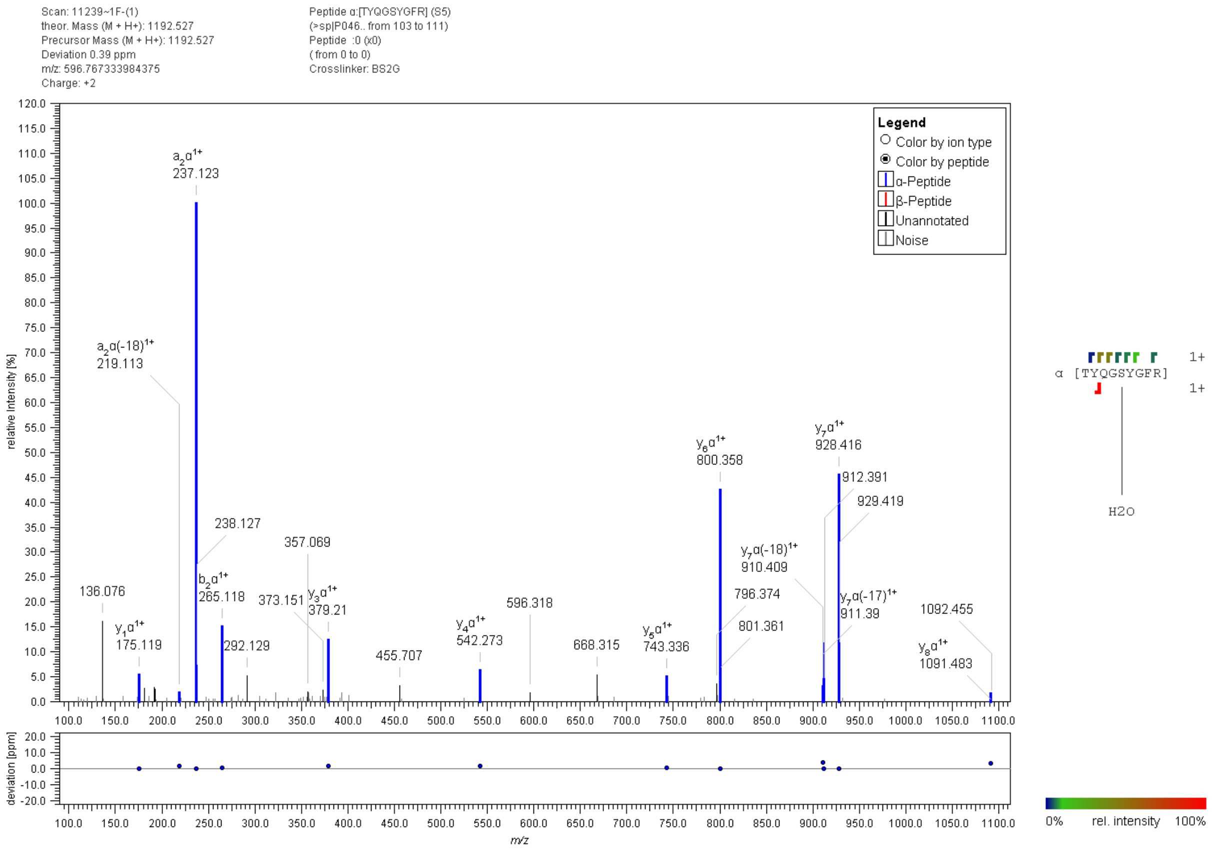


**Figure S29:** Fragment ion mass spectrum of a p53 "dead-end" cross-link with $BS^2G$. Signal assignment was performed with MeroX 2.0.1.7 (39).

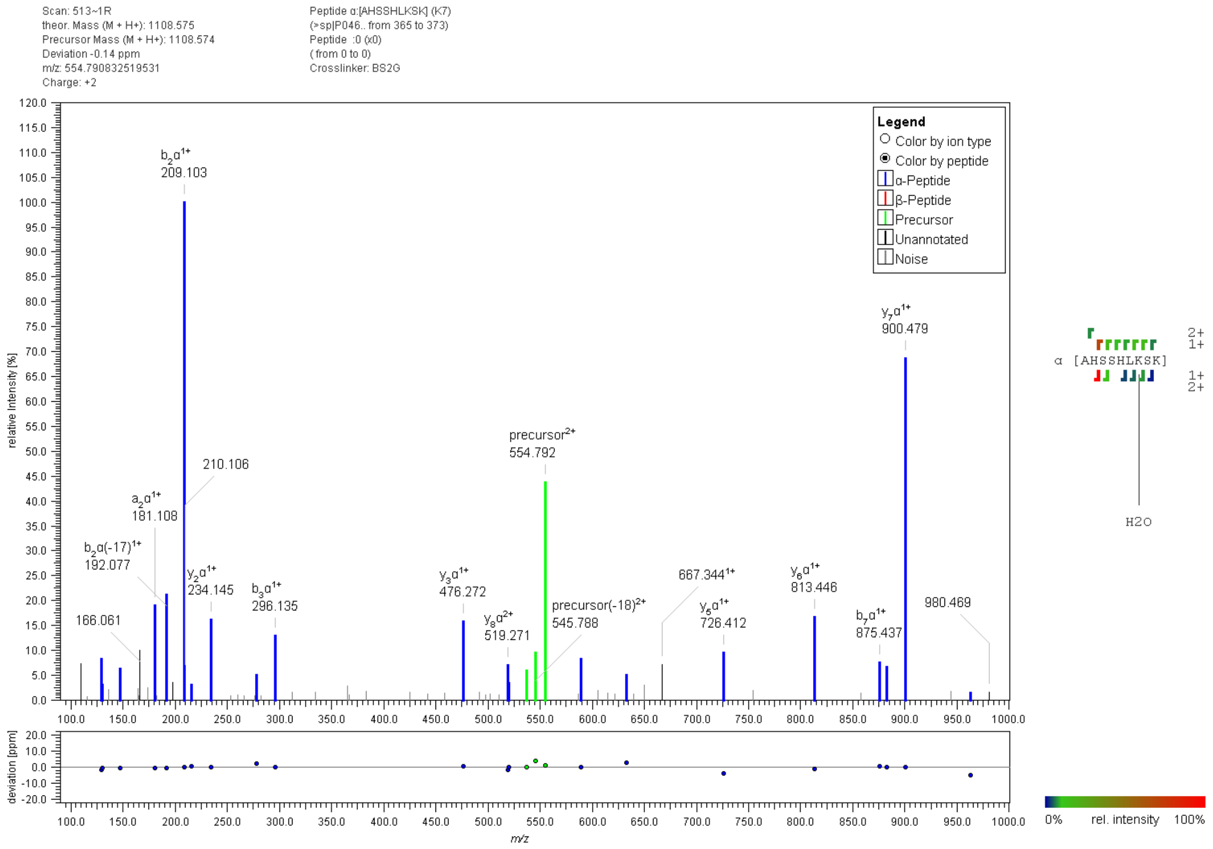


**Figure S30:** Fragment ion mass spectrum of a p53 "dead-end" cross-link with $BS^2G$. Signal assignment was performed with MeroX 2.0.1.7 (39).

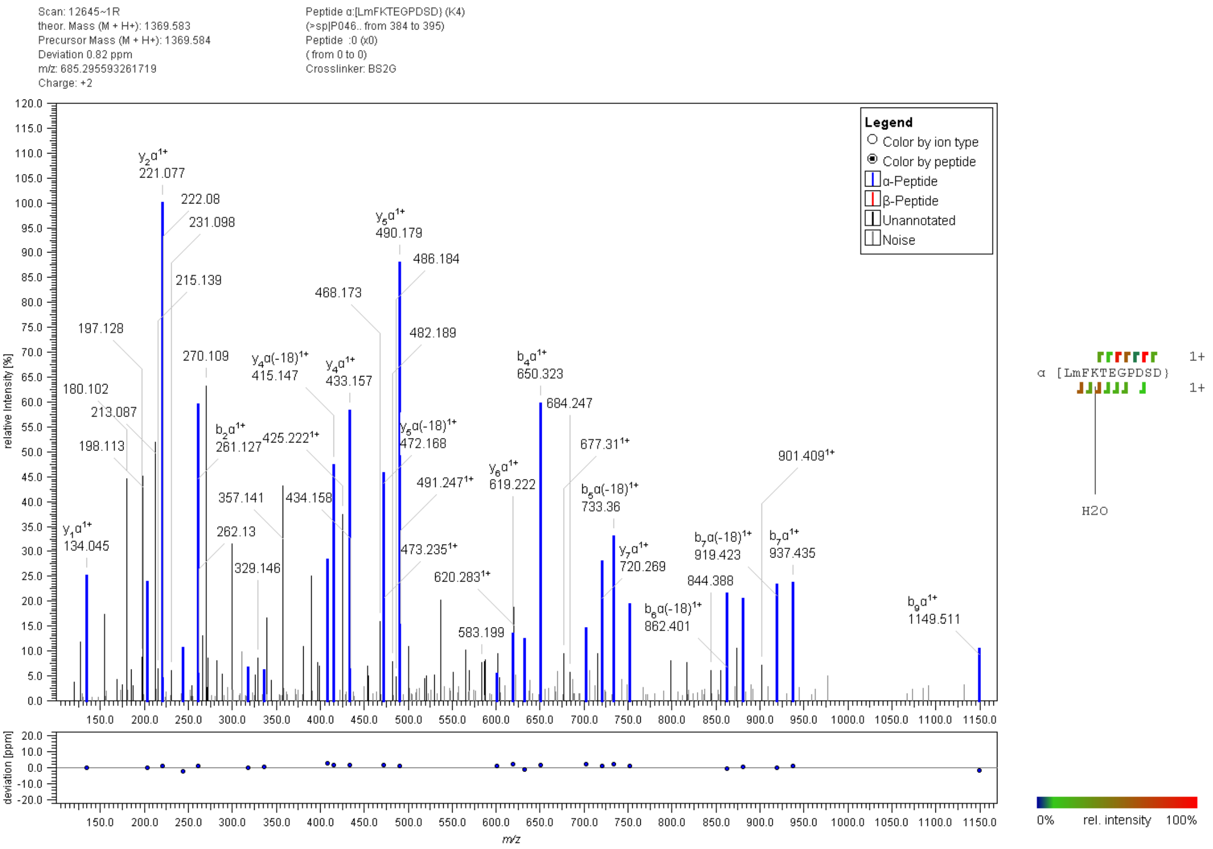


**Figure S31:** Fragment ion mass spectrum of a p53 "dead-end" cross-link with BS$^2$G. Signal assignment was performed with MeroX 2.0.1.7 (39).

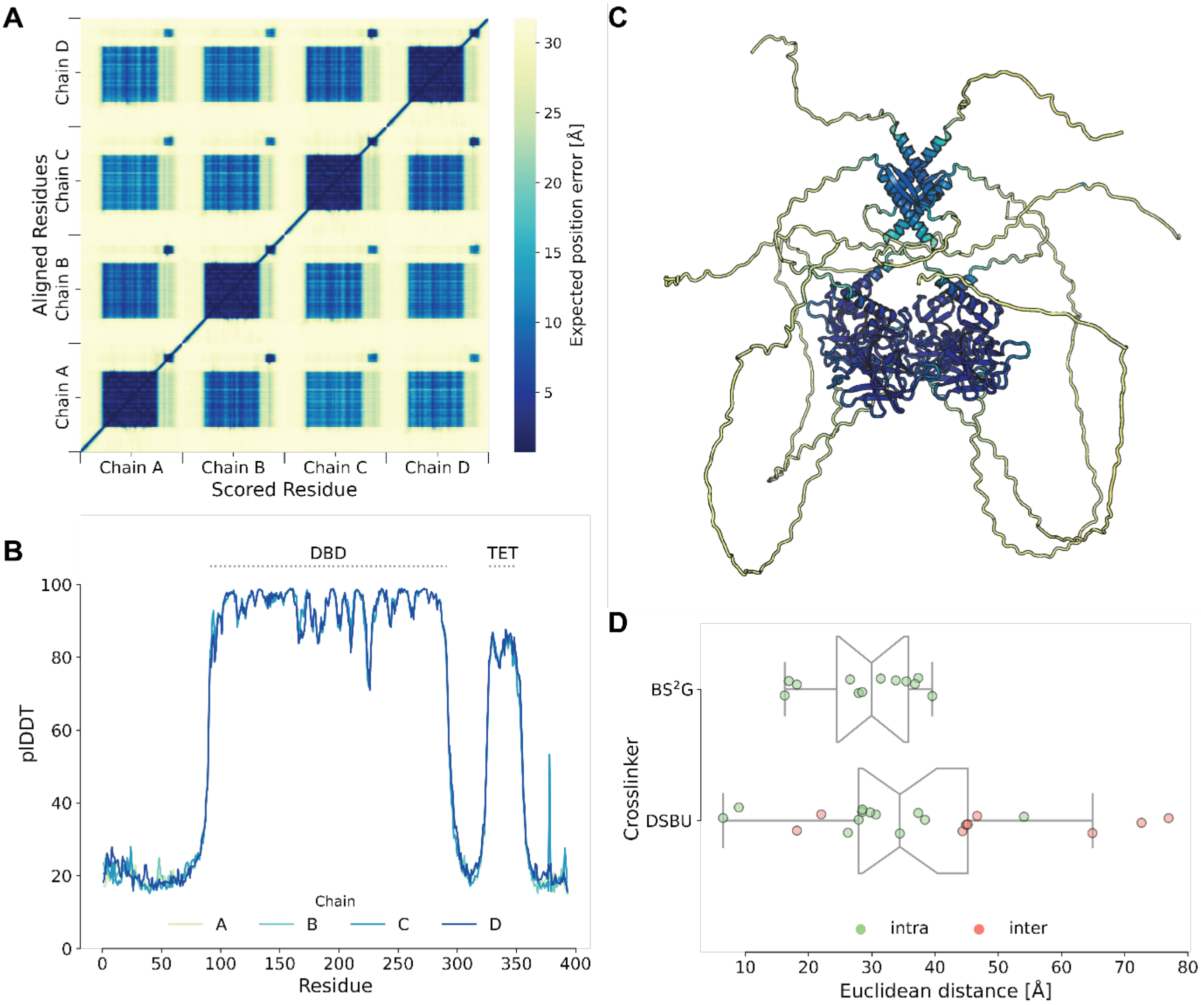


**Figure S32.** AlphaFold2 prediction of tetrameric p53. **(A)** Heatmap of the predicted aligned error of tetrameric p53. Only the structure of the (DBD; blue) has a high score. (B) AlphaFold2 confidence scores (plDDT) of each residue. Only the DBD and the TET are predicted with high confidence, indicating an ordered structure. (C) AlphaFold2 predicted tetrameric p53 atomic model in cartoon representation; coloring is based on plDDT scoring. (D) Distance distribution of XL-MS data; the shortest distance of each cross-link is plotted.

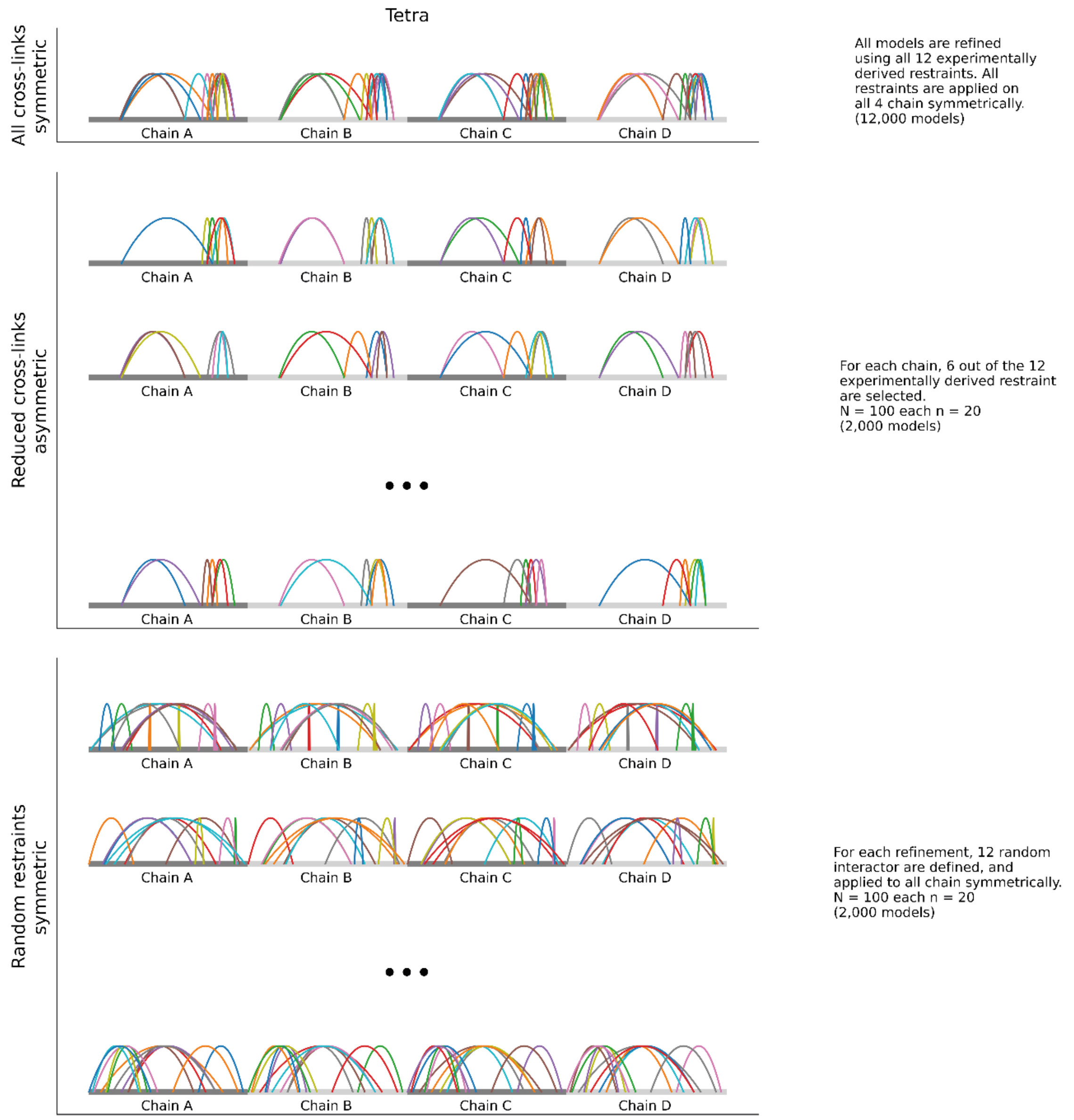


**Figure S33.** Refinement strategies. 'All Restraints' applies the same force symmetrically on each chain of p53, generating 12,000 models. In 'Reduced Restraints', half of the restraints were omitted for each chain individually, resulting in an asymmetric refinement. 100 unique refinements were performed, generating 20 models each. In 'Random Restraints', 12 restraints were created and applied symmetrically to each p53 chain. 100 unique refinements were performed, generating 20 models each.

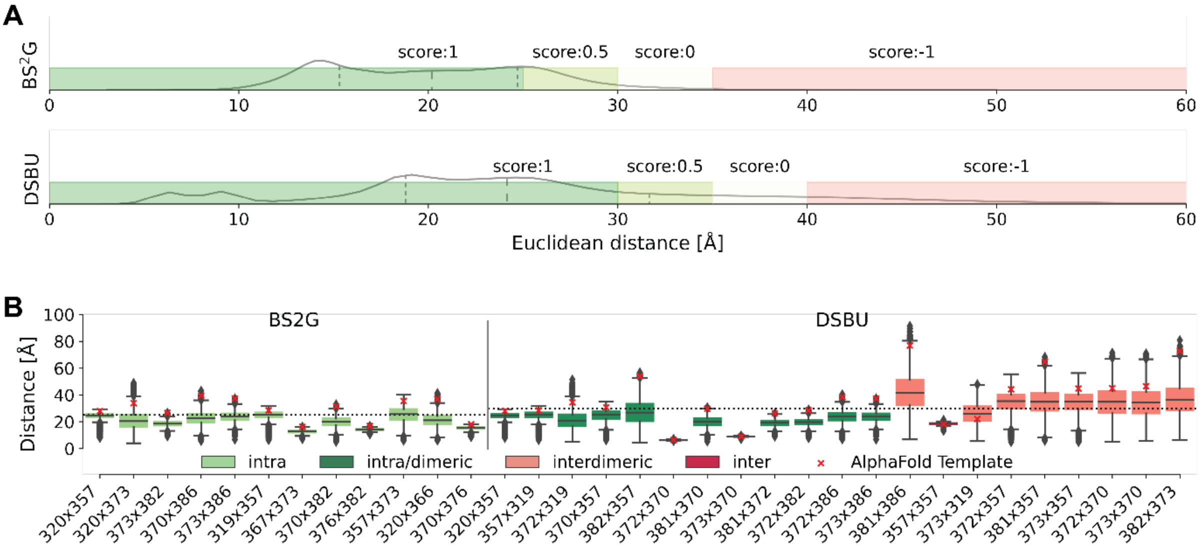


**Figure S34.** Scoring of XL-MS-derived distance restraints. (A) The plots indicate the distance distributions of all refined p53 models. Based on these distributions, a maximum Euclidean Cα-Cα distance threshold was set to 25 Å ($BS^2G$) and 30 Å (DSBU), with a 5-Å stepwise scoring to account for flexibility. Scoring is indicated by color coding: dark green (perfect score 1), light green (score 0.5), white (score 0), and light red (score -1). (B) Distance distributions per cross-link pair for all models. Dotted lines indicate the maximum Cα-Cα distances allowed to achieve a perfect score. Red crosses indicate Cα-Cα distances in the initial AlphaFold2 model (template).

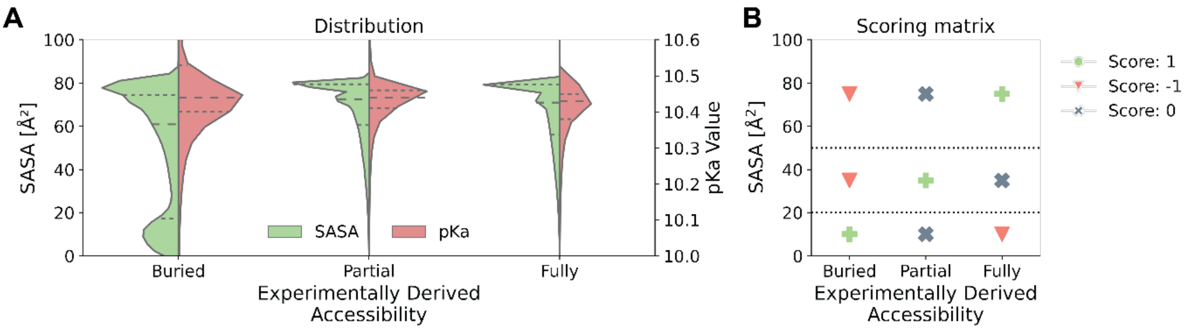


**Figure S35.** Scoring of labeling efficiency. (A) SASA and $pK_a$ values of labeled lysine residues in p53, plotted as function of footprinting data (experimentally derived accessibility). Although $pK_a$ values are unaltered there seems to be a population of lysine residues that are not accessible to labeling, as is visible from changing SASA values. (B) Scoring matrix of footprinting data.

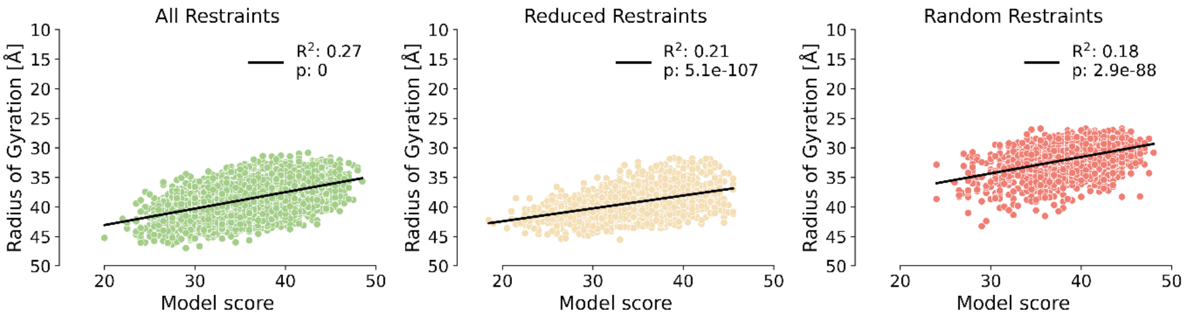


**Figure S36.** XL-MS and footprinting data are in agreement with a compact structure of p53 tetramer. In all refinement strategies, a significant correlation ($p < 1x10^{-50}$) is observed, with more compact structures giving a better scoring.

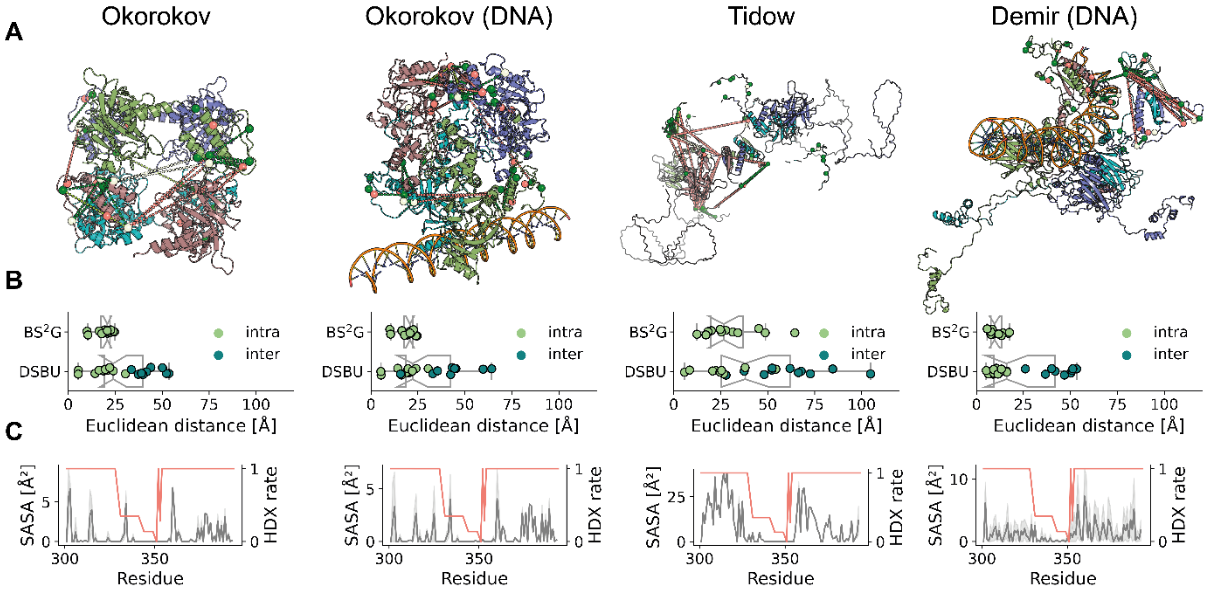


**Figure S37.** Existing models of p53 higher-order structures poorly explain our XL-MS and footprinting data. (A) Atomic tetrameric p53 models according to Okorokov *et al*, 2006 (+/-DNA) (19); Tidow *et al*, 2007 (-DNA) (20); Demir *et al*, 2017 (+DNA) (25). Cross-links are indicated as dashed lines between Cα atoms of residues involved; residues labeled in fottprinting experiments are indicated by colored spheres. Color code corresponds to Figure S34. (B) Distance distributions of BS$^2$G and DBSU cross-links show distance violations between p53 dimers in the existing models of p53 tetramer; intra: cross-links within one p53 monomer, inter: cross-links between p53 dimers (38). (C) Existing models show a high correlation between SASA (grey) and HDX (orange). The disordered and accessible nature of p53's IDRs is reflected by high SASA and HDX rates, while the structured TET (residues 326-356) exhibits low SASA and HDX rates.

**Table S1:** $BS^2G$ and DSBU cross-linking data with Cα-Cα distances for all models generated in this study (file TableS1.csv).

**Table S2:** SASA scoring of labeled lysine residues for all models generated in this study (file TableS2.csv).

**Table S3:** $BS^2G$ cross-links of p53 (+/- RE-DNA); cross-linked residues are underlined.

| Cross-link sequence | Cross-linked residues |
|---|---|
| $^{320}$KKPLDGEYFTLQIR$^{333}$<br>$^{352}$DAQAGKEPGGSR$^{363}$ | K320-K357 |
| $^{159}$AMAIYKQSQHMTEVVR$^{174}$<br>$^{292}$KGEPHHELPPGSTK$^{305}$ | Y163-K292 |
| $^{320}$KKPLDGEYFTLQIR$^{333}$<br>$^{373}$KGQSTSR$^{379}$ | K320-K373 |
| $^{373}$KGQSTSR$^{379}$<br>$^{382}$KLMFK$^{386}$ | K373-K382 |
| $^{383}$LMFKTEGPDSD$^{394}$<br>$^{364}$AHSSHLKSK$^{372}$ | K370-K386 |
| $^{383}$LMFKTEGPDSD$^{394}$<br>$^{373}$KGQSTSR$^{379}$ | K373-K386 |
| $^{307}$ALPNNTSSSPQPKK$^{320}$<br>$^{352}$DAQAGKEPGGSR$^{363}$ | K319-K357 |
| $^{364}$AHSSHLKSK$^{372}$<br>$^{373}$KGQSTSR$^{379}$ | S367-K373 |
| $^{364}$AHSSHLK$^{370}$(SK)<br>$^{382}$KLMFK$^{386}$ | K370-K382 |
| $^{382}$KLMFK$^{386}$<br>$^{374}$GQSTSR$^{379}$ | S376-K382 |
| $^{352}$DAQAGKEPGGSR$^{363}$<br>$^{373}$KGQSTSR$^{379}$ | K357-K373 |
| $^{364}$AHSSHLKSK$^{373}$<br>$^{374}$GQSTSR$^{379}$ | K370-S376 |

**Table S4:** Footprinting of lysine residues in p53 (+/- RE-DNA). Average labeling is compared between DNA-bound and DNA-free p53 and labeling ratios are calculated (presented in Figure 2); p-value is set to < 0.05.

| Labeled residue number | Avrg $DNA^-$ % labeling | Avrg $DNA^+$ % labeling | $log_2(DNA^+/DNA^-)$ | $log_{10}$(p-value) |
|---|---|---|---|---|
| K120 | 20.38 | 20.14 | -0.02 | 0.10 |
| K139 | 0.81 | 0.51 | -0.68 | 0.55 |
| K165 | 26.22 | 29.91 | 0.19 | 0.82 |
| K291 | 3.29 | 3.15 | -0.06 | 0.05 |
| K292 | 8.92 | 12.22 | 0.45 | 0.32 |
| K305 | 21.77 | 25.15 | 0.21 | 0.21 |
| K319 | 19.31 | 25.39 | 0.39 | 0.81 |
| K320 | 2.21 | 2.27 | 0.04 | 0.18 |
| K321 | 9.04 | 9.45 | 0.06 | 1.68 |
| K351 | 3.03 | 2.34 | -0.37 | 0.17 |
| K357 | 14.44 | 4.95 | -1.54 | 0.34 |
| K370 | 30.81 | 34.97 | 0.18 | 0.28 |
| K372 | 7.56 | 8.83 | 0.22 | 0.58 |
| K373 | 16.24 | 18.41 | 0.18 | 0.29 |
| K381 | 17.88 | 17.98 | 0.01 | 0.01 |
| K382 | 21.28 | 22.64 | 0.09 | 0.14 |
| K386 | 22.54 | 30.95 | 0.46 | 0.55 |
